# Buried Ion-Pairs Reshape Folding Pathways of Artificial Proteins

**DOI:** 10.64898/2026.10.10.757878

**Authors:** Jobst Liebau, Grant Kemp, Dirk Auman, Niklas Hoja, Alexander Jussupow, Mona Baumgart, Gunnar von Heijne, Ville R. I. Kaila

## Abstract

**Abstract:** Ion-pairs are central charged motifs that control the biological function of several proteins. Yet, the physical principles linked to the formation of ion-pairs during protein folding remain poorly understood. Here, we show that buried ion-pairs of *de novo*-designed helical bundle proteins reshape the folding pathways by introducing kinetic barriers that are overcome by a hydrophobic collapse linked to packing of the protein core. By combining protein design with co-translational folding studies, nuclear magnetic resonance (NMR) spectroscopy, and molecular dynamics simulations, we show that introduction of a buried ion-pair leads to a shifted folding transition that balances desolvation and electrostatic interactions within the protein and the bulk solvent. Moreover, we find that the folding pathway is sensitive to modifications in non-polar interactions and motifs that electrostatically shield the ion-pair from its surroundings. Our combined findings provide a blueprint for understanding how functional motifs affect protein folding and how mutation of charged networks alter the stability of proteins.

**Significance Statement:** Despite significant progress in machine learning-based structure prediction, we still lack a clear understanding of the folding pathways that determine how proteins fold into their exact three-dimensional structure. Here, we show that introduction of a buried ion-pair reshapes the folding pathway of *de novo*-designed protein *Maquettes*. By combining cotranslocation folding experiments of these proteins trapped within the ribosome exit tunnel with chemical unfolding experiments using nuclear magnetic resonance spectroscopy and atomistic simulations, we show that the ion-pair introduces a folding barrier linked to formation of the protein core. On a general level, our findings highlight how charged networks balance electrostatic and solvation forces in proteins, and how mutations can alter their folding landscape.

## Introduction

Ion-pairs, formed between oppositely charged amino acids, establish key electrostatic interactions that regulate the stability and biological function of several proteins (1–9). At the protein surface, ion-pairs contribute to protein stability and solubility (2–4, 10), enable the formation of specific protein complexes (11), and, when embedded within the protein core, ion-pairs can control the enzymatic and ion-transport activity (6, 12). In the respiratory Complex I, buried ion-pairs regulate proton transport across a biological membrane (5–7), whereas in the molecular chaperone heat-shock protein 90 (Hsp90), they provide gates that couple the enzymatic activity with conformational dynamics of the protein (13). Elucidating the physical properties of buried ion-pairs is thus of key importance in deriving a mechanistic understanding of complex proteins.

In contrast to solvent-exposed charged networks, buried ion-pairs can introduce significant (up to 5 kcal mol^-1^) destabilizing effects on the overall protein stability. These effects arise from the (Born) desolvation free energy, linked to moving the charged residues from a high dielectric (ε_w_=80) aqueous environment in the unfolded (U) state to the low-dielectric protein interior (ε_p_=4-10) of the folded (F) state (14–18), which are larger than the respective electrostatic interactions between the oppositely-charged sidechains of the ion-pair. Although such effects have been established for both natural and artificial proteins, it still remains unclear how the buried ion-pairs may reshape folding pathways and influence the free energy barriers during the folding process. Of particular biological importance is co-translational protein folding, in which the nascent polypeptide chain initiates the folding process inside or upon exiting the ribosome tunnel (19–22).

Designed 4-helix protein “*Maquettes”* provide minimal synthetic model systems to study biological function without the complex biological surroundings of natural systems, and the possible “evolutionary baggage” of natural protein sequences (23–28). Despite their simple coiled-coil design, various *Maquette* proteins have been shown to perform a wide range of enzymatic functions, such as ion-transport (27), oxidoreductase activity (24, 25), stereoselective cyclopropanation (28), and they have been employed to study light-capturing effects (26), oxygen-binding (23, 24), as well as buried charged networks (18).

Here, we probe how buried ion-pairs affect the co-translational folding trajectory of artificial helical bundle proteins by studying the effect of how interactions amongst charged residues within the protein core affect the folding process. We probe how the buried ion-pairs introduced into an artificial *Maquette* protein affect the folding processes by combining force profile analysis (FPA) (19–22) with chemical unfolding experiments, nuclear magnetic resonance (NMR) spectroscopy, and atomistic molecular dynamics simulations. Based on the combined data, we suggest a model for how buried ion-pairs affect the folding of *de novo*-designed helical bundle proteins, with implications for how complex charged networks are created in natural proteins.

## Results

To address the folding principles of the artificial *Maquette* proteins, we expressed and purified three artificial helix bundle constructs based on previous designs (18) – a fully packed four helical bundle with a hydrophobic packed core (HC), a construct containing a buried ion-pair (IP) introduced at position E17/K72, as well as a charge-buffered ion-pair (IP’), in which the ion-pair is shielded by introducing a hydrogen-bonding N69 residue (IP-V69N), next to K72 (**Fig. 1A**). Experimental structures of the folded state of HC and IP show that the coiled-coil structure forms helix-loop-helix motifs, with helix 1 adjacent to helices 2 and 4 (**Fig. 1A**). While NMR structures of HC and IP have been determined (18, 26), no structure for IP’ exists. We therefore modelled an IP’ structure based on the NMR-model of IP by introducing the corresponding mutation *in silico* (**Fig. 1A**). To obtain molecular insights into the folding principles, we studied the co-translational folding of the constructs using force profile analysis (FPA), and their chemical unfolding using CD and NMR spectroscopy, in combination with atomistic molecular dynamics (MD) simulations.

**Figure 1.**
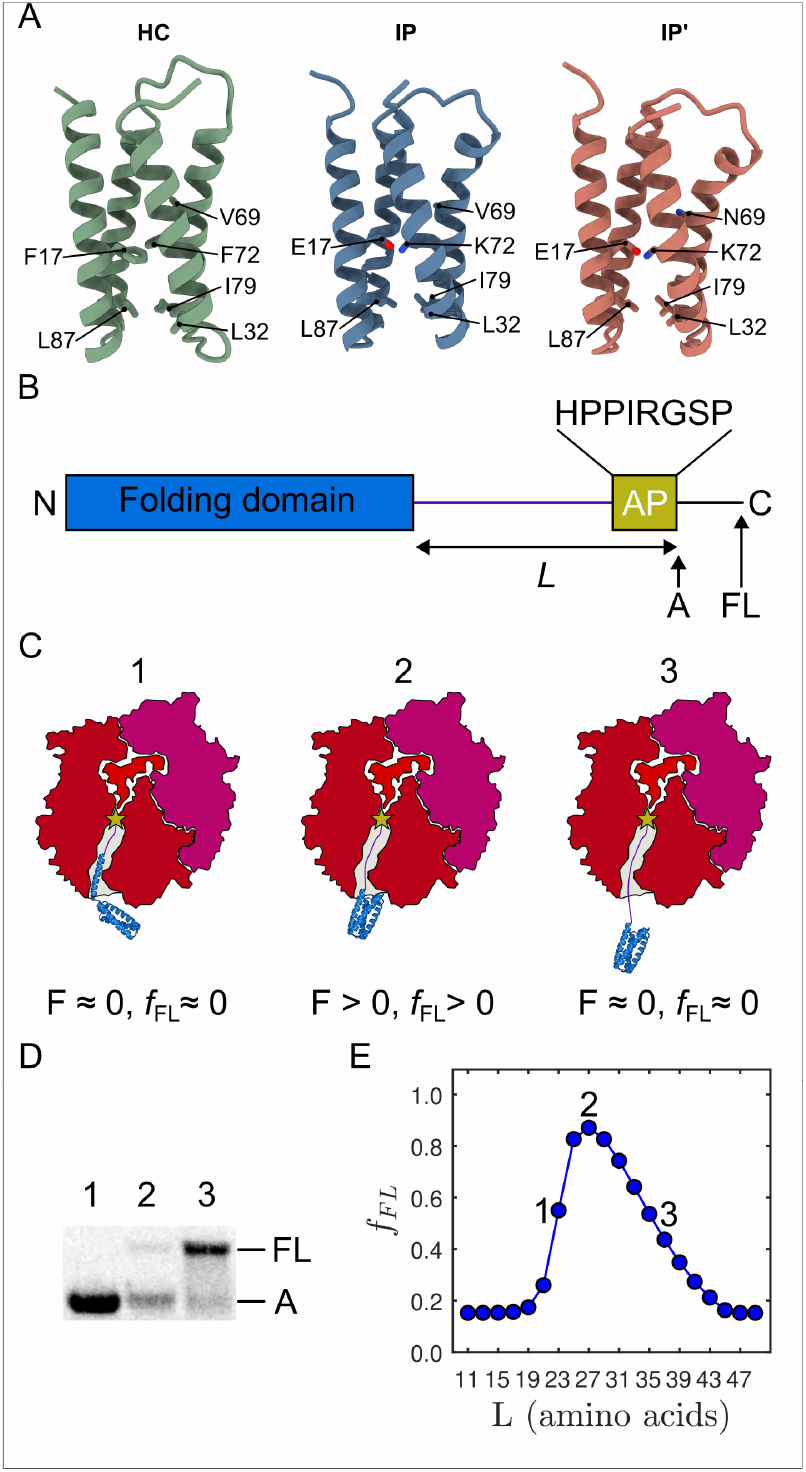
Structure of protein *Maquettes* and overview of the force profile analysis (FPA).

(A)Structure of HC (green, PDB ID: 5TGW), IP (blue, PDB ID: 6Z35), and an *in silico* model of IP’ (red). The ion-pairs (E17/K72) in IP and IP’, and analogous non-polar residues (F17/F72) in HC are shown as sticks. (**B**) Principles of the FPA: An N-terminal folding domain (blue) is fused to a linker region (purple) of variable length *L*, an arrest peptide (AP), and a C-terminal extension sequence that allows the separation of the arrested (*A*) from the full-length (*FL*) protein by SDS-PAGE. (**C**) Schematic representation of different scenarios during an FPA experiment: Protein folding can induce a force in the nascent chain that allows the interactions of a force-sensitive AP with the ribosome to be overcome and for the translation to continue. 1) For a short *L*, all residues required for folding are not yet available when the ribosome reaches the AP (asterisk), leading to a low force generated on the nascent chain (*f*_FL_≈0), and high yield of arrested product. 2) Intermediate *L* enable folding, and the ribosome translates the AP, while the folding generates a force on the nascent chain, leading to the production of *FL* protein (*f*_FL_>0). 3) For large *L*, the folding domain has already folded before the AP is translated by the ribosome, leading to low generated force, and little production of *FL* product (*F*≈0, *f*_FL_≈0). (**D**) An autoradiograph of the SDS-PAGE gel, showing the separation of *A* and *FL* protein. Lane 1 - arrest control with the AP immediately followed by a stop codon, resulting in a product that is equivalent in size to *A*. Lane 2 - experimental sample showing both *A* and *FL* products. Lane 3 – full-length control with a mutant, non-functional AP that generates a product equivalent in size to *FL*. (**E**) Idealized force profile, showing *f*_FL_ as a function of *L* based on the model in SI Appendix, Fig. S1-S3. *L_onset_* (1, see **C**) and *L_offset_* (3, see **C**) are defined as the half-maximal onset and offset of the force peak, respectively, and *L_max_* (2, see **C**) as the maximum of the peak.

### The force profile analysis (FPA) assay

Translational arrest peptides (APs), initially discovered as regulators of the translation process (29– 32), are short segments of polypeptide that, immediately upon synthesis on the ribosome, stall protein translation by forming tight interactions with the ribosome exit tunnel (ET) in the vicinity of the peptidyl-transferase center (PTC) (33). Because the stalling efficiency of an AP is sensitive to pulling forces acting on the nascent chain (19, 34, 35), APs can be engineered to sense the forces generated by co-translational folding of a target protein (21, 36), here the co-translational folding of the synthetic *Maquette* proteins with the introduced charge networks (IP, IP’) in relation to the same model with a packed hydrophobic core (HC).

In FPA, the co-translational protein folding process is studied by designing a series of protein constructs, in which the target protein is fused, via a variable-length linker of length *L* residues, to an AP followed by a short C-terminal tail **(Fig. 1B)**. As illustrated in **Fig. 1C** and **D** (*left*), in constructs with short linkers (*i.e*., low *L* values), in which the target protein has not yet reached a location in the ET that is wide enough to be compatible with folding when the ribosome reaches the last codon of the AP, little force will be generated on the AP and the arrested (*A*) form of the protein will predominate, as also suggested by a two-state model of the cotranslation folding process (see SI Appendix, **Fig. S1-S3**). In contrast, at a critical linker length (**Fig. 1C** and **D,** middle), the target protein reaches a location in the ET, where it can start to fold at the same time as the ribosome reaches the last codon in the AP; the folding reaction generates a pulling force on the AP that weakens its stalling efficiency and results in increased synthesis of full-length (*FL*) protein. Finally, at long linker lengths, for which folding is already completed when the ribosome reaches the end of the AP, little pulling force is generated on the AP, again resulting in mostly arrested (*A*) protein product (**Fig. 1C** and **D,** right; SI Appendix, **Fig. S1**). Thus, by plotting the fraction of full-length translated protein (*f_FL_*)

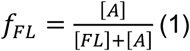

as determined from SDS-PAGE gels against the linker length (*L*) (**Fig. 1D**), we constructed a force profile (FP, **Fig. 1E**), in which the folding transition will appear as a peak. Moreover, under conditions where *f_FL_* linearly depends on the pulling force, the total work *W* exerted along the translated distance can be calculated by integrating *f_FL_* over the folding peak,

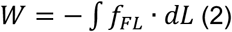

See SI Appendix for a qualitative model that relates *f*_FL_ to the free energy of folding (Δ*G_fold_*).

### Buried ion-pairs affect the co-translational folding of Maquette proteins

For construct HC with the packed core, the FP shows a sharp onset of the folding transition with *L_onset_* ≈ 24 residues (**Fig. 2A**, green curve), a peak maximum at *L_max_* ≈ 27 residues, followed by a gradual drop in *f*_FL_ with *L_end_* ≈ 30 residues. Interestingly, the ion-pair introduced into construct IP leads to a strongly perturbed FP: the folding transition takes place at a later stage as compare to HC (*L_onset_* ≈ 26 residues, *L_max_* ≈ 29 residues, *L_end_* ≈ 31 residues), and the overall exerted work *W*, estimated from the area under the curve, is reduced to ∼40% relative to HC **(Fig. 2B**, blue bar). These profiles are consistent with prior circular dichroism (CD) chemical unfolding experiments (**Fig. 2C–E**, *cf*. also Ref. 18) as well as NMR experiments (see below), indicating that the buried ion-pair destabilizes the overall protein fold by a few kcal mol^-1^ (SI Appendix, **Fig. S4A**).

**Figure 2.**
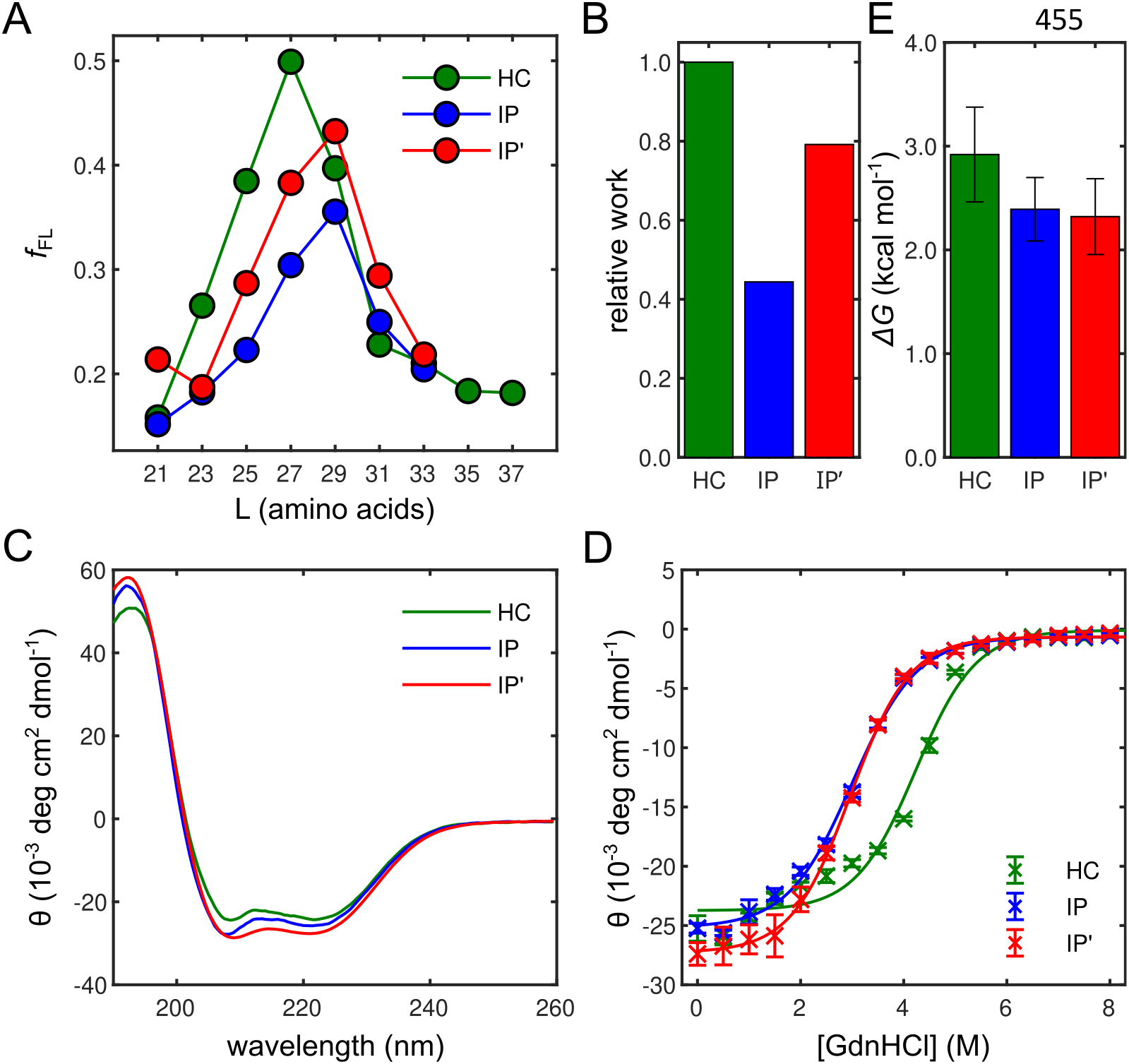
Co-translational folding of protein *Maquettes* studied by force profile analysis and chemical unfolding experiments.

(A) FPA of the protein *Maquettes* HC (green), IP (blue), and IP’ (red). The folding transitions shift from HC (*L_onset_* = 24 residues) to IP (*L_onset_* = 26 residues) due to introduction of an ion-pair within the hydrophobic core. IP’ has a similar folding transition as IP, but the folding is initiated earlier (*L_onset_ =* 25 residues), indicating a shift in the folding co-operativity. **(B)** Relative work *W* exerted along the translated distance normalized to the HC data. **(C)** CD spectra, **(D)** GdnHCl unfolding experiments, and **(E)** free energy derived from CD chemical unfolding experiments for HC, IP, and IP’.

Based on protein design and statistical analysis of natural proteins, it was suggested (18) that buried ion-pairs can be stabilized by amphipathic charge-shielding residues that form both hydrogen-bonded contacts with the ion-pair and non-polar contacts with the protein core. Introduction of such a charge-shielding asparagine residue near the ion-pair in construct IP’ (F17E/F72K/V69N) leads to folding peak that is in-between that of HC and IP (**Fig. 2A**, red curve). As compared to IP, IP’ has a larger Δ*G_fold_* and a higher folding co-operativity, as indicated by an increase of the exerted work *W* (**Fig. 2B**) and by chemical unfolding experiments (18) (**Fig. 2C-E,** SI Appendix, **Fig. S4A**), These findings suggest that the free energy gained from packing the non-polar core is partially used to compensate for the energetic penalty introduced by the buried ionpair, and that the charge-shielding asparagine residue indeed significantly stabilizes the ion-pair.

To further probe how the non-polar interactions affect the folding transition, we replaced bulky leucine and isoleucine residues with smaller alanine residues (L32A, I79A, L87A) to stepwise destabilize the hydrophobic packing for HC, IP, and IP’. As expected, chemical unfolding experiments of these destabilized variants, suggest that each substitution (1A - 3A) decreases the protein stability by around 0.5 kcal mol^-1^ (SI Appendix, **Fig. S4A**). The perturbations of the non-polar interactions are clearly visible also in the force profiles, with the amplitude of the folding peak and the exerted work *W* gradually reducing with increasing numbers of substitutions (SI Appendix, **Fig. S4C-H**). In parallel, the peak maxima shift towards higher *L* values, supporting the notion that a significant amount of non-polar interactions must form in order to surpass the solvation effects linked to formation of the buried ion-pair (see below).

### In vitro folding of Maquette proteins

To further probe the role of the ion-pairs in destabilizing the hydrophobic protein core during the folding pathway, we next conducted chemical unfolding experiments probed by nuclear magnetic resonance (NMR) spectroscopy. In the absence of urea, all three constructs display disperse heteronuclear-single quantum coherence (HSQC) spectra, indicating that they adopt well-folded structures (**Fig. 3A**). In increasing urea concentrations, the unfolding gives rise to less disperse spectra, as all amide backbone groups experience similar chemical environments in the unfolded state. For IP and IP’, these transitions qualitatively occur around 4 – 5 M urea concentrations, while the HC is substantially more stable and displays disperse spectra even at ∼8 M urea (**Fig. 3A**), consistent with the CD experiments (see above).

**Figure 3.**
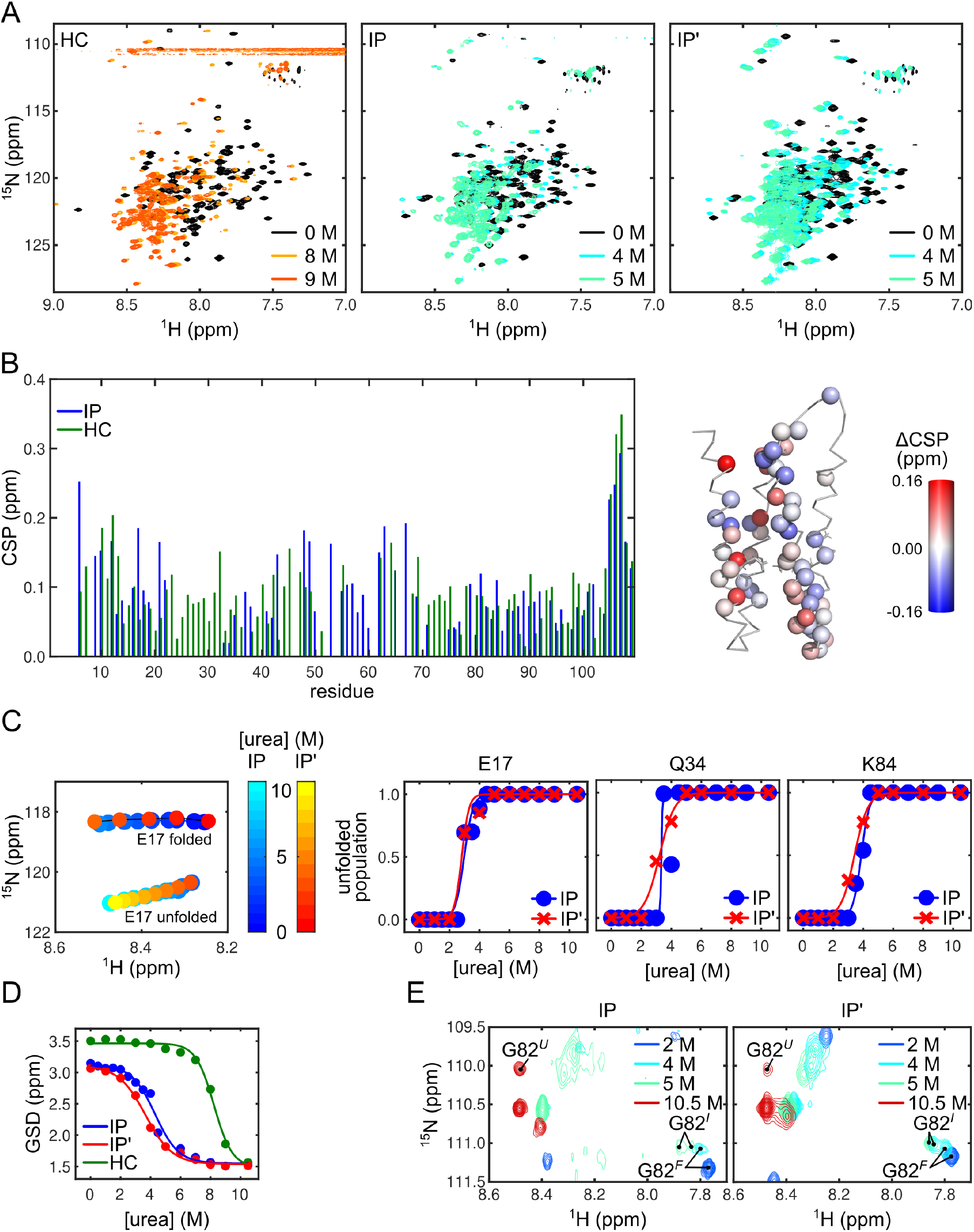
Unfolding of protein *Maquettes* studied by NMR spectroscopy.

(**A**) ^15^N-HSQC of HC, IP, and IP’ at various urea concentrations. (**B**) CSPs of ^15^N-HSQC obtained for 0 M and 3 M of urea (*left*) and differences in CSP (ΔCSP = CSP^IP^ – CSP^HC^) mapped onto the NMR structure of IP (PDB ID: 6Z35). ΔCSP < 0 indicates residues in IP that are less surface-exposed than in HC, and *vice versa* for ΔCSP > 0. (**C**) Unfolding analysis of IP and IP’. Residues are traced in ^15^N-HSQCs at increasing urea concentrations from the folded state, and decreasing populations of the unfolded state (top *left*). Populations of the unfolded and folded states in IP and IP’ at increasing urea concentrations (top *right* and bottom) for E17, Q34, and K84. Eq. 9 (see SI Appendix, *Extended Methods*) is fitted to the data, with parameters listed in SI Appendix, Table S1A. (**D**) Global spectral dispersion (GSD) for IP, IP’, and HC at increasing urea concentrations (see SI Appendix, Table S3, for parameters and fitting). (**E**) ^15^N-HSQC for IP (*top*) and IP’ (*bottom*) of the glycine region at various concentrations of urea. *F* – folded conformation; *I* – intermediate conformation; and *U* – unfolded conformation.

To obtain more quantitative insights into the unfolding mechanism, we assigned the HSQC spectra of the folded (F) and unfolded (U) states for all constructs, partially using previous assignments of the F state (Refs. 18, 26, *cf*. also BMRB ID: 30185 (HC), BMRB ID: 34518 (IP)). For HC, we are unable to assign the U state, possibly because even at 10 M urea the F and U states coexist (SI Appendix, **Fig. S5**), whereas for the unfolded IP and IP’, we can assign up to 80% and 65% of residues, respectively (SI Appendix, **Fig. S6**).

To detect possible structural differences, we evaluated chemical shift perturbations (CSPs) between conditions without urea and upon 3 M urea addition that resulted in the unfolding transition (**Fig. 3B**). In these conditions, we observe both positive and negative CSPs between IP and HC (ΔCSP^IP/HC^) (**Fig. 3B,** SI Appendix, **Fig. S7**), suggesting that the introduced ion-pair impacts the packing of helices during the folding process, qualitatively consistent with the folded structures of IP and HC (SI Appendix, **Fig. S8**). At the folding transition, the ion-pair exchanges between *open* and *closed* conformation, in which the packing differences localize to helices 1 and 2 (*closed* conformation) or to helices 3 and 4 (*open* conformation), for which we observe the strongest ΔCSP^IP/HC^ (**Fig. 3B,** SI Appendix, **Fig. S7**), supporting conformational flexibility at the folding transition, consistent with structural differences between HC and IP. In contrast, the CSP differences experienced by IP relative to IP’ (ΔCSP^IP/IP’^) are minor, indicating overall similar structures (SI Appendix, **Fig. S7**). These observations are further corroborated by a direct comparison of the HSQCs, which, despite only two mutations, differ substantially between folded IP and folded HC, while the spectra of the folded IP and IP’ are similar (SI Appendix, **Fig. S9**).

By assignment of the unfolded and folded conformations of IP and IP’, we could trace resonances of the U state with decreasing urea concentrations, as well as, resonances of the F state with increasing urea concentrations. Since the proteins unfold and refold on a slow NMR timescale, resonances of the U and F states coexist at intermediate urea concentrations, allowing us to quantify populations of U and F spectra (**Fig. 3C**). In total, we traced 10 resonances for IP and IP’ (**Fig. 3C,** SI Appendix, **Fig. S10**), while discarding traces that overlap or lack assignments for spectra in the 0-10 M urea titration range. For all traceable resonances, the folding transitions are similar for IP and IP’ (SI Appendix, **Table S2**), with an overall unfolding transition occurring at 3.6±0.5 M urea for IP and 3.1±0.4 M for IP’, assuming a two-step transition.

To probe the global folding process, instead of individual residues that can be assigned under all conditions, we defined a global spectral dispersion (GSD) of the HSQC spectra as a measure to characterize the reduction in overall spectral dispersion in the ^1^H dimension of the HSQC. We define the GSD as the sum of spread of HSQC resonances from their center of mass, weighted by the distance of each resonance from the center of mass in the ^1^H dimension (see SI Appendix, *Extended Methods*), and calculated as a function of increasing urea concentration (**Fig. 3D**). The GSD suggest that HC has a steeper folding transition (dGSD/d[urea]∼-1.65×10^-3^ ppm M^-1^ with a midpoint at 8.7 M urea) than IP (dGSD/d[urea]∼-1.28×10^-3^ ppm M^-1^, midpoint at 4.3 M urea), whilst IP has a steeper folding transition relative to IP’ (-1.11×10^-3^ ppm M^-1^ with the midpoint at 3.7 M urea). The GSD analysis suggests that IP’ stabilizes intermediates along the folding pathway as compared to IP, in line with a shifted co-operativity of the folding transition. In this regard, the two-state unfolding process is most likely an over-simplification of the model, as indicated by resonances of G82 that show the co-existence of a folding intermediate for both IP and IP’ (**Fig. 3E**). For HC, the non-linear chemical shift perturbations of G82 (SI Appendix, **Fig. S11**) indicate that also HC adopts a folding intermediate, which, in contrast to IP and IP’, are in fast exchange with the folded conformation, further supporting that the folding co-operativity is altered in the helical bundles with a buried ion-pair.

Taken together, our NMR experiments show that the introduction of a buried ion-pair into the non-polar core of an artificial helical bundle protein results in a shifted folding pathway. Our NMR experiments resolve intermediate states, indicative of a stepwise dissociation of the helices during the folding process. While the IP and IP’ constructs undergo a folding transition around similar urea concentration, the differences in folding co-operativity, as revealed by the global spectral dispersion and putative folding intermediates, could qualitatively reflect perturbation in the folding pathway as observed in our co-translational folding experiments.

### Exploring the molecular principles underlying folding of Maquette proteins

To derive a molecular understanding of the chemical unfolding experiments, we simulated the urea-induced unfolding process by atomistic molecular dynamics (MD) simulations in different urea concentrations (0-10 M) and temperatures (300-400 K) using the a99SB-disp force field (37), which provides an improved balance between folded and unfolded regions as compared to other biomolecular force fields. Notably, the constructs sample partially unfolded conformations at high temperatures and urea concentrations on the accessible simulation timescale (144 μs cumulative simulations, 4 μs per simulation condition), suggesting that our simulations primarily capture early and intermediate unfolding events (**Fig. 4**). We further analyzed the unfolding trajectories by a principal component analysis (PCA) combining all data, clustered independently for each construct to identify representative conformational ensembles.

**Figure 4.**
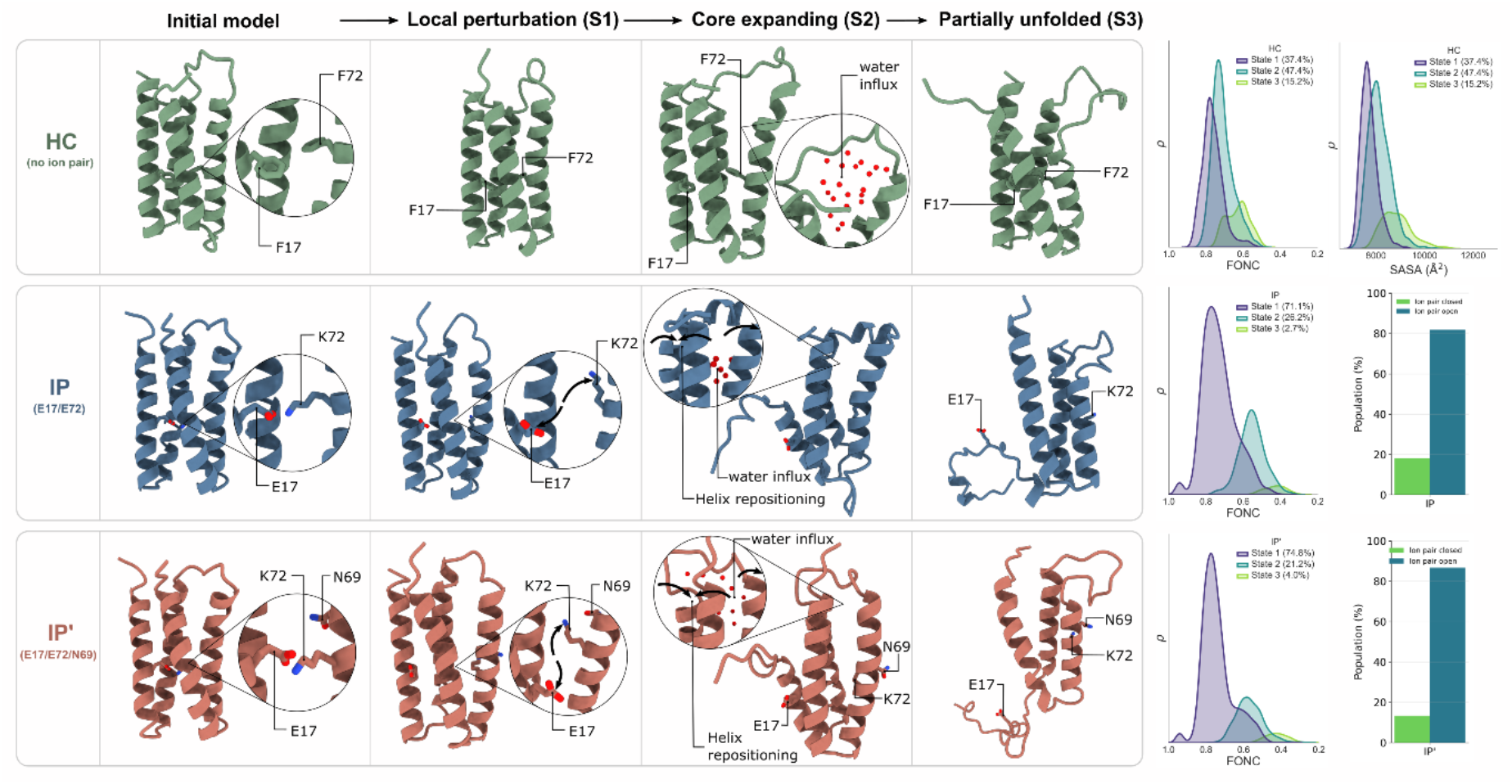
Conformational ensembles identified from unfolding simulations of protein *Maquettes*.

Principal component and clustering analysis based on MD simulation of protein Maquettes (HC, IP, IP’) to identify representative conformations along the unfolding pathway. Structures are organized according to increasing structural disruption from native-like conformations through locally perturbed (S1), core-expanded (S2), and partially unfolded states (S3), representing putative unfolding intermediates. Insets highlight structural rearrangements, including ion-pair opening, helix repositioning, and water influx into the hydrophobic core. HC samples a structurally distinct conformational ensemble as compared to IP and IP’, showing local unfolding originating primarily from rearrangements around the loop region of helices 1 and 2. In contrast, the unfolding of IP and IP’ is characterized by a displacement of helix 1, following ion-pair rearrangement and core hydration. The assignment of putative unfolding intermediates is based on analysis fraction of native contacts (FONC), solvent exposed surface area (SASA), and ion-pair and hydration states.

The PCA clustering reveals recurrent partially unfolded intermediates that differ between HC and the IP and IP’ variants, consistent with our FPA and NMR data showing that the ion-pair reshapes the folding pathway (**Fig. 4**). For HC, we observe an unfolding process that is initiated by expansion of the hydrophobic core, accompanied by increased water penetration into the protein interior. This process couples to rearrangements of the loop region between helices 2 and 3, and leads to local destabilization and progressive expansion of the protein core, while largely maintaining the overall bundle architecture. In contrast, during early steps of unfolding transition of IP and IP’, the ion-pair opens up, and reorients toward the solvent, suggesting that destabilization of the charged interaction network precedes larger structural rearrangements. The ion-pair opening is also accompanied by a local rearrangement of the bundles that result in dissociation of helices 2 and 4. These rearrangements coincide with a partial displacement of helix 1, increasing the hydration of the hydrophobic core, and expansion of the loop region between helices 2–3. Dissociation of helix 1 from the bundle results in a progressive rearrangement of helices 2 and 4, giving rise to partially unfolded conformations with increased solvent exposure and extended loop regions, with qualitatively similar unfolding pathways for IP and IP’. In this regard, IP’ samples a somewhat more expanded and dynamic conformational ensembles under some simulation conditions, although these differences remain small relative to the common unfolding features shared by both ion-pair variants.

## Discussion

We have studied here how ion-pairs embedded within the non-polar core of artificial 4α-helical bundle proteins affect the protein folding pathways by cotranslation folding assays in combination with spectroscopic studies and molecular simulations of the unfolding process. Our findings can be rationalized in terms of physical effects underlying the forces that govern the folding process. The hydrophobic interactions within the folding core and the hydration free energy linked to embedding of charged residues are expected to increase with the size of the folding intermediate, with contributions arising both from the increasing surface area and volume of the folding protein fragments, linked to the surface tension and van-der-Waals packing energy (38). In contrast, the dielectric constant of the emerging protein core gradually decreases along the folding co-ordinate (λ), and embedding the charged residues into the protein interior thus results in an enhanced electrostatic interaction between the ion-pair, which counteracts an increased (Born) desolvation free energy for transferring the charged residues from the bulk solvent (ε_w_=80) to the protein interior (ε_p_(λ)). These effects thus disfavors embedding the charges as well as the ion-pair dipole into the folding protein core with an increasing λ. In this regard, our MD simulations suggested that the embedding of the ion-pair is coupled to specific changes in its local conformational environment: folded-like conformations predominantly showed a restricted, buried ion-pair geometry, whereas the partially unfolded conformations showed disruption of the ion-pair that led to solvent-exposed configurations, with water molecules solvating the charged sidechains. This increased conformational heterogeneity could result in entropic effects, where both the local residue as well as the water re-arrangement influence the overall charge embedding process.

Although our simulations did not capture a continuous unfolding trajectory, we combined simulation data across different temperatures and urea concentrations, to sample partially unfolded conformational states that reveal structural rearrangements and suggest putative early steps during the unfolding process. In particular, HC predominantly samples intermediates characterized by progressive hydration and expansion of the bundle core, whereas IP and IP’ populate ensembles associated with ion-pair opening, local structural rearrangements, and increased solvent penetration into the protein core. These observations suggest that incorporation of a buried ion-pair alters the sequence of structural rearrangements along the folding pathway, rather than simply changing the overall stability of the folded state. In this regard, our combined findings across co-translational folding assays, NMR experiments, and molecular simulations, show that a buried ion-pair reshapes the folding landscape of the artificial protein constructs, with the buried ion-pair constructs populating distinct intermediate states relative to the protein with a well-packed non-polar protein core. These findings suggest that incorporation of buried charges modifies the folding pathway by introducing competing electrostatic and desolvation constraints that must be compensated during formation of the protein core.

Taken together, the folding pathway balance between favorable non-polar packing and electrostatic stabilization with the unfavorable solvation free energy that results in delicate enthalpy-entropy compensation effects. Although the exact microscopic contributions remain unclear, we can qualitatively rationalize the folding pathways along a 2D-free energy surface, with the abscissa reflecting the progression of hydrophobic core formation, whereas the ordinate represents a reaction co-ordinate associated with core desolvation and ion-pair re-organization (**Fig. 5**). Assuming a gradual lowering of ε_p_ along the folding co-ordinate λ, a minimum free energy folding pathway is expected along the concerted ion-pair- and non-polar packing co-ordinate, suggesting that the ion-pair forms at a later stage along the folding pathway relative to HC, containing only a non-polar core region.

**Figure 5.**
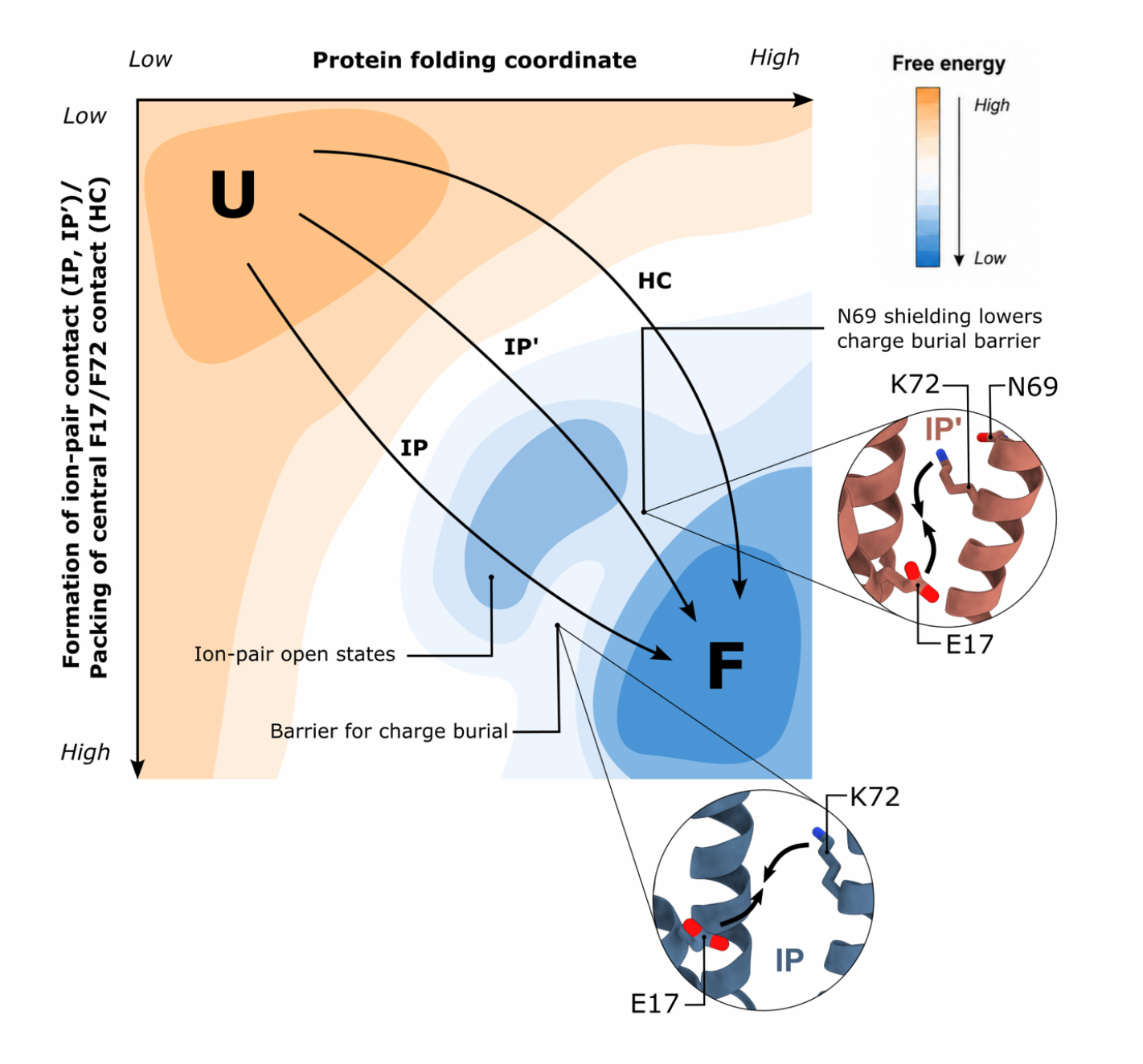
Schematic folding landscape of the protein *Maquettes*.

Schematic 2D free-energy landscape, illustrating alternative folding pathways of the protein *Maquettes* with a non-polar hydrophobic core (HC), and buried ion-pair (IP), and charge-shielded ion-pair (IP’). The horizontal co-ordinate represents the packing of the hydrophobic core, whereas the vertical co-ordinate represents formation of the ion-pair (for IP, IP’) or packing of the central F17/F72 (for HC) contact, with unfolded (U) and folded (F) states marked. HC follows a direct downhill pathway dominated by a hydrophobic collapse, followed by formation of the central F17/F72 at late stages of the folding processes. IP and IP’ fold by forming a partially assembled hydrophobic core, with a solvent-exposed open-ion pair state. The folding then proceeds via complete ion-pair burial and dehydration of the protein core. In the IP variant, this step is associated with a larger desolvation and charge-burial barrier relative to IP’, where Asn69 partially shields the desolvation penalty, and allows for more packing of the non-polar core.

Despite the recent AI revolution in predicting the structure of folded proteins (39–41), our findings emphasize the intricate and still poorly understood molecular principles that determine the folding landscape of both natural and artificial proteins. Our study shows how the introduction of a single ion-pair into the protein core reshapes the folding landscape, and illustrate the delicate balance between non-polar packing effects and electrostatic interactions. Although the studied artificial proteins show a conformational landscape distinct from natural proteins, they highlight thermodynamic and kinetic principles that also natural proteins must optimize during the folding and assembly of extended functional networks of charged residues, as observed in various proteins, ranging from molecular chaperones to membrane-bound proteins responsible for energy transduction.

## Materials and Methods

### Cloning

A full list of protein sequences is included in SI Appendix, **Table S1**. Constructs for biophysical characterization were synthesized by GenScript (Piscataway, USA) with an N-terminal hexahistidine and TEV protease cleavage site (ENLYFQS), and cloned into a pET21a vector. The initial constructs for Force Profile Analysis (FPA) were also synthesized by GenScript with the same N-terminal tag and cleavage site and cloned into a pET21a vector with the following sequences fused to the C-terminus of *Maquette*: a flexible tetrapeptide (SGSG), a random coli linker sequence of variable length derived from *E. coli* LepB (more detail below), an HA-tag (YPYDVPDYA), the MsSecM(sup1) arrest sequence (HPPIRGSP), and a 23 amino acid-long sequence to allow the resolution of the arrested (AR) and full-length (FL) products by SDS-PAGE (see **Fig. 1D**). The LepB linker sequence varied in length from 0 (*L*=21) to 40 (*L*=61) amino acids at steps of two amino acids. The linker length, L, for each construct is the number of amino acids from the flexible SGSG tetrapeptide to the C-terminal proline of the arrest peptide. The minimal (*L*=21) sequence contains no added LepB linker sequence. Additional intermediate linker lengths were created using partially overlapping reverse PCR to truncate the linker sequence. The linker was always truncated from the N-terminus such that the sequence closest to the ribosome (immediately upstream of the HA tag) would not change to avoid linker length-specific interactions with the ribosomal exit tunnel. This linker strategy has been used in several previous publications (19–22). Introduction of substitutions in the *Maquette* sequence within the FPA constructs was accomplished either using Gibson assembly (42) to introduce sequences synthesized by Genscript with the FPA construct backbone or by site-directed mutagenesis (SI Appendix, **Table S1**).

### Protein expression and purification

The C-terminally His_6_-tagged proteins were overexpressed in *E. coli* BL21 (DE3) for 4 h at 37°C. The expression was induced with 0.5 mM isopropyl β-D-1-thiogalactopyranoside (IPTG) at OD_600_ 0.7, after which the cells were disrupted by sonication and the lysate was further purified by Ni-NTA affinity chromatography followed by size exclusion chromatography (SEC) using a preparative Superdex 75pg HiLoad 16/600 column (GE Healthcare). The His_6_-tag was cleaved by a Tobacco Etch Virus (TEV) protease, and removed by reverse IMAC.

### Circular dichroism spectroscopy

A Jasco J-715 system combined with a co-oling-heating unit was used for measuring CD spectra and chemical unfolding experiments. The measurements were performed at 20°C using a 1-mm-path quartz cuvette and a protein concentration of 10 µM. Chemical unfolding with guanidine-hyrochloride (GdnHCl) up to 8 M was performed to study the protein stability, and the folding free energy was computed from linear least square fits of -*RT* ln *K* extrapolated to 0 M GdnHCl. The equilibrium constant *K* defined as [U]/[F] was obtained from the fraction of denatured protein *f*_D_ = (*y*-*y*_F_)/(*y*_U_-*y*_F_), where *y* is the CD raw signal, and *y*_F_ and *y*_U_ corresponds to the native and denaturated protein. Error bars were computed from the standard deviation of triplicate measurements.

### Force Profile Analysis

Force Profile Analysis (FPA) was carried out as described before (19–22). Briefly, the *Maquette* open reading from the pET21a vectors described above was amplified by PCR using T7 sequencing primers (forward: 5′-CCCGCGAAATTAATACGACTCACTATAGGG-3’; and reverse: 5’-GCTAGTTATTGCTCAGCGG-3’) and purified using a GeneJet PCR purification kit (Thermo Scientific). 0.4 µL of the PCR product was added to a 5 µL (final volume) PURExpress (NEB) reaction containing ∼1 µCi of ^35^S-methionine (Perkin Elmer) and incubated at 37°C for 20 min with shaking at 600 rpm in a 1.5 mL microcentrifuge tube. The reaction was stopped by adding 5 µL of ice-cold 10% trichloroacetic acid (TCA) and allowed to rest on ice for at least 30 min. The total precipitated protein sample was pelleted in a microcentrifuge at 20,000 g for 5 min at 4°C. The supernatant was aspirated away and the pellet was resuspended in SDS-PAGE sample buffer (50 mM Tris pH 6.8, 0.1% SDS, 10% glycerol, 5 mM 2-mercaptoethanol) containing 0.6 mg/mL RNAseA. The radiolabelled arrested (AR) and full-length (FL) proteins were separated by SDS-PAGE (Invitrogen, 12% NuPAGE precast Bis-Tris gels with MES running buffer). The radiolabelled bands were detected by autoradiography and an image generated in a Fuji FLA-9000 scanner. The band pixel density was determined from the TIFF image in Fiji software. These band densities were fit to Gaussian curves for quantification by the in-house EasyQuant software. The fraction of full-length protein (*f_FL_*) was calculated as the amount of arrested protein, ([AR]), divided by the total protein, ([FL] + [AR]) using Eq. 1.

The force profile was obtained by plotting the *f_FL_* against the linker length, *L*. The force profiles were plotted in EasyQuant and the total area under the peak estimated by a sum of Gaussians. Unless otherwise noted all chemicals were purchased from Sigma-Aldrich and molecular biology enzymes from NEB.

### Nuclear Magnetic Resonance (NMR) spectroscopy

Heteronuclear ^1^H,^15^N-TROSY-HSQC spectra were recorded using ^15^N-labelled protein at 25 °C in 50 mM sodium phosphate buffer, pH 7.5 with 0-10 M urea and with 10% D_2_O for frequency-locking placed in a 5 mm diameter NMR tube (Wilmad 541-PP-8) on a Bruker Avance III spectrometer operating at a ^1^H Larmor frequency of 700 MHz (14.09 T) using a TCI triple-resonance cryoprobe. Spectra were processed using the NMRPipe/NMRDraw software suite (version 11.7) (43) and spectra were traced in Sparky v3.115 (44). For each protein construct, a total of 11 ^1^H,^15^N-TROSY-HSQC spectra (0-10 M urea, in 1M increments) were recorded and for IP spectra were additionally recorded at 0.5, 1.5, 2.5, 3.5, and 4.5 M urea. See Supplementary Material for a detailed discussion of the NMR unfolding analysis. Backbone amide resonance assignments of the constructs HC and IP in the folded state were mapped from the HSQC assignments based on Ref 18 and assignments from IP were transferred to IP’, for which only minimal spectral perturbations were observed aside from residues directly neighboring the mutation site (SI Appendix, **Fig. S9B**). Backbone assignments for IP and IP’ in the unfolded state were obtained from 3D CBCACONH, CBCANH, HNCACO, HNCA, HNCOCA, HNCO experiments in phosphate-buffered protein solutions containing 10 M urea. These experiments were performed on Bruker Avance III spectrometers operating at a ^1^H Larmor frequency of 700 MHz (14.09 T) and 900 MHz (22.31 T) using a TCI triple-resonance cryoprobe. ^1^H, ^15^N chemical shifts were assigned using Cara (version 1.9.1.7) (45).

### Molecular simulations

Molecular dynamics simulations were performed using GROMACS 2021.3 (46) employing the amber99SB-disp force field (37). The protein structures were placed in cubic simulation boxes (8 × 8 × 8 nm^3^), solvated using the TIP4P water model, and neutralized by adding Na^+^ and Cl^−^ ions to a final concentration of 150 mM (SI Appendix, **Fig. S12**). Systems were energy minimized, followed by *NVT* and *NPT* equilibration prior to production simulations. Unfolding simulations were performed at urea concentrations ranging from 0–10 M and temperatures between 300–400 K using a 2 fs integration timestep, resulting in 144 μs cumulative simulation trajectories. Long-range electrostatic interactions were treated using the particle mesh Ewald (PME) method. Trajectories were analyzed using the fraction of native contacts (FONC), solvent accessible surface area (SASA), and in-house scripts (SI Appendix, **Fig. S13-S15**). Visualization was performed using MDAnalysis (47, 48), VMD (49) and UCSF ChimeraX (50).

### Principal component analysis

Trajectories from all simulation conditions (144 μs, 0–10 M urea, 300–400 K) were combined and analyzed by principal component analysis (PCA) using Cα co-ordinates after alignment to the initial structure. All constructs were projected into a common PCA space using the first five principal components (PC1–PC5) (SI Appendix, **Fig. S16**).

Conformations were subsequently clustered separately for each construct in the shared PCA space into five clusters to identify representative conformational ensembles (SI Appendix, **Fig. S17-S19**). Clusters displaying similar structural features were subsequently combined into representative unfolding intermediate ensembles.

## Acknowledgments

We thank Cecilia Persson at the Swedish NMR Centre for technical assistance with the NMR experiments. This work received funding from the Knut and Alice Wallenberg Foundation (2019.0251 and 2024.0220 to V.R.I.K, 2022.0241 to G.v.H.), the Göran Gustafsson Foundation for Research in Natural Sciences and Medicine (to V.R.I.K.), the Novo Nordisk Fund (NNF18OC0032828 to G.v.H), and the Swedish Research Council (2025-04607, 2020-04081 to V.R.I.K. and 2020-03238 to G.v.H), and the European Research Council (ERC, bioenergetic, to V.R.I.K.) This project has received funding from the European Research Council (ERC) under the European Union’s Horizon Europe research and innovation programme (grant agreement No 101265268). Computational resources were provided by the National Academic Infrastructure for Supercomputing in Sweden (NAISS 2026/1-56, NAISS 2025/1-33, 2024/1-28, 2023/1-31) for computational resources.

