## Supplementary material for "Buried Ion-Pairs Reshape Folding Pathways of Artificial Proteins": SI Appendix

**This PDF file includes:**

**Extended Methods**
- Qualitative model of force generation during co-translational folding
- NMR Analysis

**Supplementary Figures**
- Fig. S1. Co-translational folding of a protein inside the ribosome exit tunnel
- Fig. S2. Folding free energy in the ribosomal exit tunnel.
- Fig. S3. Force profiles for various AP constructs.
- Fig. S4. The effect of decreasing hydrophobicity on the stability and folding of the maquette constructs.
- Fig. S5. $^{15}$N-HSQC of HC at 10.5 M urea concentration.
- Fig. S6. Assignment of unfolded state of IP and IP'.
- Fig. S7. Chemical shift perturbation of protein Maquettes during chemical unfolding experiments.
- Fig. S8. Structural analysis of HC and IP.
- Fig. S9. HSCQ spectra of HC, IP, and IP'.
- Fig. S10. Assignment of residues during the unfolding transition in NMR.
- Fig. S11. Resonance of the folded conformation monitored by G82 of HC at increasing urea concentrations.
- Fig. S12. Molecular dynamics simulation setup.
- Fig. S13. Fraction of native contacts (FONC) during unfolding simulations.
- Fig. S14. Solvent-accessible surface area (SASA) during unfolding simulations.
- Fig. S15. Distance distributions, monitoring core rearrangements during unfolding simulations.
- Fig. S16. Principal component analysis of conformational ensembles during unfolding simulations.
- Fig. S17. PCA derived cluster analysis of HC unfolding intermediates.
- Fig. S18. PCA derived cluster analysis of IP unfolding intermediates.
- Fig. S19. PCA derived cluster analysis of IP' unfolding intermediates.

**Supplementary Tables**
- Table S1. Sequences of studied Maquette constructs.
- Table S2. Unfolding analysis of IP and IP'.
- Table S3. Parameters for the global spectral dispersion analysis of IP, IP', and HC.

**Supplementary References**

### Extended Methods

#### Qualitative model of force generation during co-translational folding

To qualitatively rationalize shifts in the FPA profiles, we derive a simple force-generation model. To this end, we consider the ribosome exit tunnel (ET) to be composed of a narrow part in which the protein domain (PD) cannot fold, and a wider part in which there is enough room for the protein to fold (**Fig. S1**). The PD is attached to an arrest peptide (AP) by an unstructured linker of equilibrium length $x_e$ (proportional to the number of residues in the linker *N*). Folding of the PD is initiated when its C-terminal end is in location $x = x_c$ in the ET. We treat the linker as an entropic spring, with spring constant $K_p$, where the energy $E_S$ stored in spring is,

$$E_S(x) = \frac{1}{2}K_p(x - x_e)^2 \quad (1)$$

and the force from the spring on the AP is,

$$F_S(x) = K_p(x - x_e) \quad (2)$$

The folding free energy of the isolated PD is $\Delta G_{U\to F}$ and the PD can fold only when $x \geq x_c$, hence, for given $x_e$ and $x_c$ (set by the number of residues in the linker and the length of the ET, respectively), the energy of the linker+PD system varies with the instantaneous location, *x*, of the C-terminal end of the PD as shown in Fig. S1. For simplicity, we only consider the free energy of the unfolded (*U*) state at $x = x_e$, and of the folded state (*F*) at $x = x_c$ (**Fig. S1**), and put $G_U = 0$, and $G_F = E_S(x_c) + \Delta G_{U\to F}$, where $\Delta G_{U\to F}$ is the folding free energy of the isolated PD. The total free energy of folding $\Delta G_{fold}$ with a linker of equilibrium length $x_e$ (**Fig. S2**) is,

$$\Delta G_{fold} = \frac{1}{2}K_p(x_c - x_e)^2 + \Delta G_{U\to F} \quad (3)$$

The equilibrium constant for the $U \leftrightharpoons F$ transition for with a linker of equilibrium length $x_e$ is,

$$K_{U-F} = \frac{[F]}{[U]} = \exp\left(\frac{-\Delta G_{fold}}{RT}\right) \quad (4)$$

and the fraction of folded PD, $f_{fold}$, with a linker of equilibrium length $x_e$ is,

$$f_{fold}(x_e) = \frac{K_{U-F}}{1+K_{U-F}} \quad (5)$$

We further assume that only the folded form of the PD exerts a force on the AP, hence the average force $\langle F\rangle$ exerted on the AP with a linker of equilibrium length $x_e$ is,

$$\langle F\rangle = f_{fold}(x_e)F_s(x_c) = \frac{\exp\left(\frac{-\Delta G_{fold}}{RT}\right)}{1+\exp\left(\frac{-\Delta G_{fold}}{RT}\right)}K_p(x_c - x_e), \quad (6)$$

Note that (6) holds only for $x_e \leq x_c$, and that $\langle F\rangle = 0$ for $x_e > x_c$.

We use the model presented in Ref. 1 to convert from $\langle F\rangle$ to fraction full-length protein ($f_{FL}$) measured in the FPA experiment,

$$k_R = k_0 \exp\left(\frac{\langle F\rangle \Delta x^{\ddagger}}{k_B T}\right) \quad (7)$$

where $x^{\ddagger}$ is the distance to the transition state and $k_0$ is the rate constant for the AP released under force $\langle F\rangle$ according to Bell's model (2),

$$f_{FL} = 1 - \exp(-k_R \Delta t) \quad (8)$$

where $\Delta t$ is the incubation time obtained from the PURE *in vitro* translation system.

This simplified model makes it possible to convert the mean force $\langle F \rangle$ (in pN) acting on the AP to $f_{FL}$. A typical example, using parameter values for the SecM(*Ec*) AP (*cf*. Ref. 1), is shown in **Fig. S3A**, and an example of a stronger AP with $k_0$ = 0.2*$k_0$[SecM(*Ec*)] shown in **Fig. S3B**. Note that $x_{max}$ does not change between the two examples.

**NMR Analysis**

Assigned peaks were mapped along increasing urea conditions in both unfolding (0→10 M urea) and folding (10→0 M urea) direction. Residues were identified based on 1) assignment in both the 0 M and 10 M spectra, and assignable from either direction, and in all urea concentrations (**Figure 3C, Fig. S10**); 2) assignment in both the 0 M and 10 M spectra, and assignable for some intermediate conditions, but with one or more of the intermediate conditions unassigned, *e.g.*, due to the highly crowded 2D spectrum with many overlapping peaks or peak broadening due to intermediate exchange dynamics; and 3) assignment in either 0 M or the 10 M spectrum, but not both. For these conditions, the peak was traced until unobservable, which was taken as the end point of the titration. Peak coordinate lists were obtained from Sparky and analyzed in Matlab v. 2022b, employing in-house scripts. To quantify unfolding for the residues in category 1 and 2, peaks were integrated in Sparky and the populations of U ($p_U$) and F ($p_F$) state in dependence of urea concentration were calculated as $p_U = I_U / (I_U + I_F)$, where $I_{U/F}$ is the integral of the U/F state resonance. At low urea concentrations all protein is folded (no U resonances detectable), while at intermediate concentrations both U and F resonance are observed. At high urea concentrations, only U exists for IP and IP' (no F resonances detectable). The folding transition can be characterized by a sigmoidal folding model assuming two states, for which the unfolded population in dependence on the urea concentration [U] is computed as,

$$p_U([U]) = M + A\frac{\exp(S([U]-[U]_{mid}))}{1+\exp(S([U]-[U]_{mid}))} \quad (9)$$

for which $[U]_{mid}$ is the mid-point urea concentration, at which F and U co-exist to 50%, $S$ is the slope of the sigmoidal function at $[U]_{mid}$, $A$ is its amplitude and $M$ is the offset. For data converted to populations, as in the case of individually traced resonances, $M$ = 0 and $A$ = 1. However, for the GSD (see below), $M$ and $A$ depend on the initial ([U] = 0 M) and final dispersion ([U] = 10 M) of the spectra and may adopt other values.

The general spectral dispersion (GSD) was obtained by calculating the center of mass (CoM) of the spectrum. The distance of each coordinate was then weighted by its intensity and distance from the CoM in the $^1$H dimension was calculated, summed up over all coordinates and normalized by the sum of all intensities,

$$\text{GSD} = \frac{\sum_y \sum_x \sqrt{(x-x_{CoM})^2} I(x,y)}{\sum_y \sum_x I(x,y)} \quad (10)$$

where $y$ is the $^{15}$N dimension, $x$ is the $^1$H dimension, $x_{CoM}$ is the CoM of the $^1$H dimension and $I(x,y)$ is the intensity for a point $(x,y)$ in the spectrum. To reduce artifacts, sidechain resonance intensities that are insensitive to secondary structure changes were set to zero, and intensities below a threshold of 0.05 times the maximum intensity were considered to be noise and set to zero. In addition, the $^{15}$N coordinate was not included in the weighting since $^{15}$N chemical shifts are substantially less sensitive to secondary structure than $H^N$ chemical shifts. Upon unfolding at increasing urea concentrations, the spectral dispersion reduces implying that the GSD decreases in a sigmoidal manner, allowing us to fit a sigmoidal two-state model to GSD (**Eq. 9**) in the same way that we fit such a model to the individually traced resonances as described above. In contrast to the individually traced resonances, the GSD allows us to globally characterize the unfolding even in the absence of complete assignments.

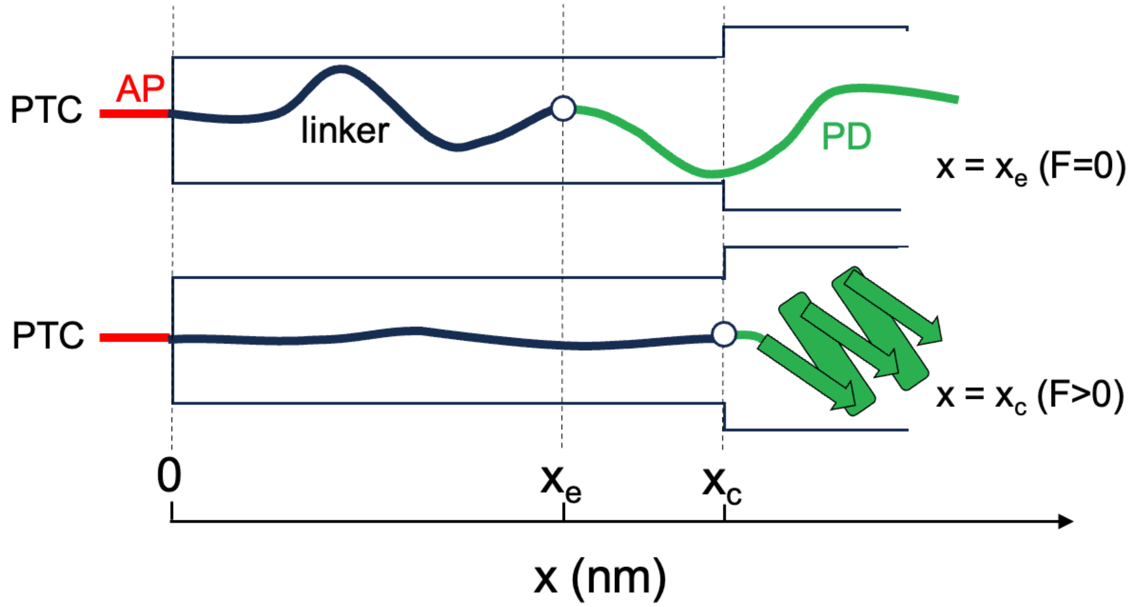


**Figure S1**. **Co-translational folding of a protein inside the ribosome exit tunnel (ET).** *x* is the instantaneous position of the C-terminal end of the protein domain (PD) in the ET (indicated by a circle). *Top*: $x_e$ is the equilibrium location of the C-terminal end of the PD in the ET, when the linker is unstretched ($x_e$ is proportional to the linker length). *Bottom*: Folding of the PD is initiated when the linker is sufficiently stretched that the C-terminal end is in location $x = x_c$ in the ET.

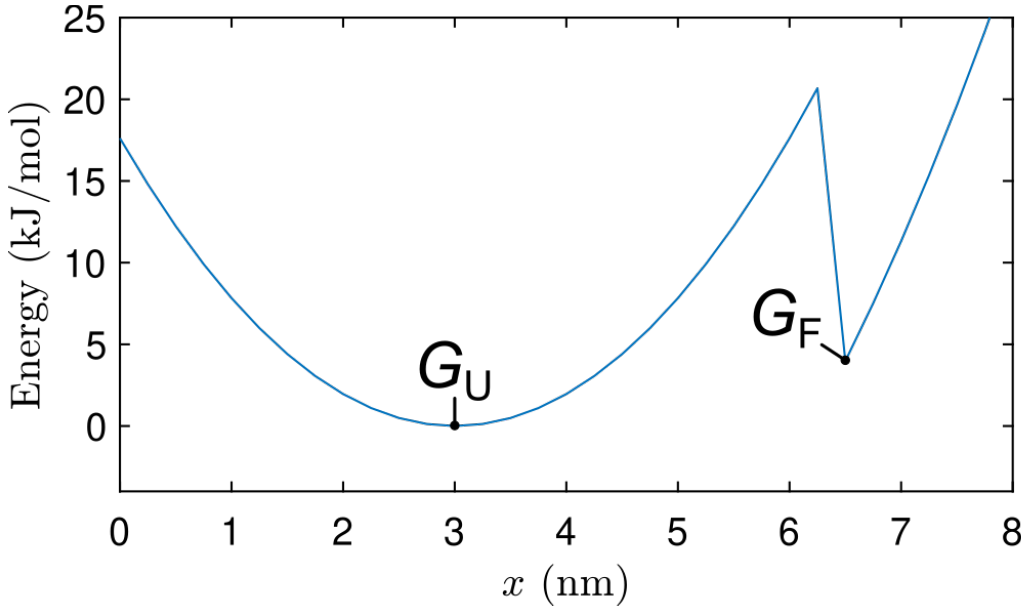


**Figure S2. Folding free energy in the ribosomal exit tunnel.** Total energy of the linker+PD as a function of the location *x* of the C-terminal end of the PD in the ET (*cf.*, Eqn. 1 in the text). The PD can fold when $x \geq x_c$, adding $\Delta G_{U\rightarrow F}$ to the total energy. The following values were assumed: $K_p$ = 6.5 pN nm$^{-1}$, $\Delta G_{U\rightarrow F}$ = -20 kJ mol$^{-1}$ (-33 pN nm), $x_e$ = 3 nm, $x_c$ = 6.5 nm. The free energies of the U and F states, $G_U$ (set equal to zero) and $G_F$, are indicated.

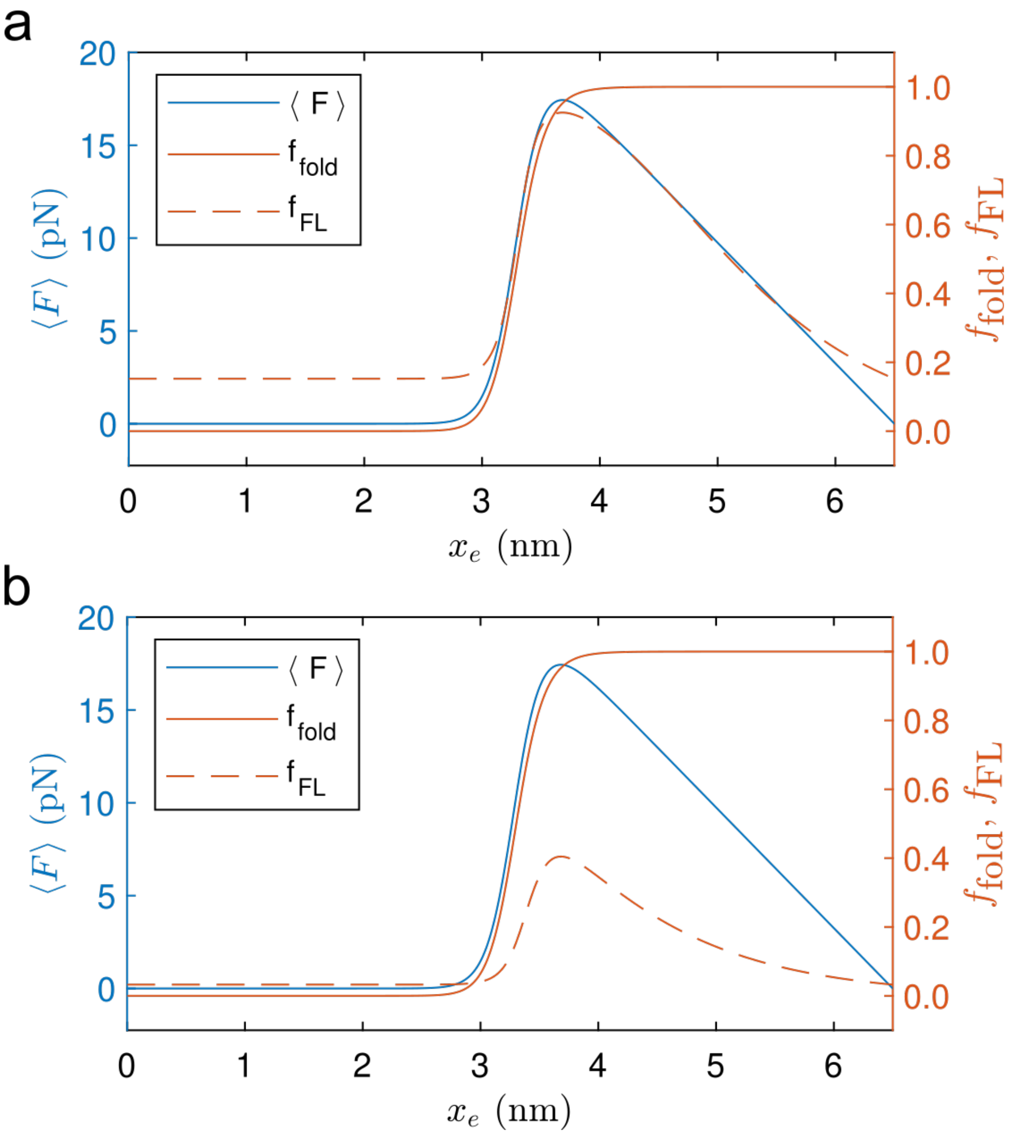


**Figure S3. Force profiles for two AP constructs: (A)** for SecM(*Ec*) AP and **(B)** for an AP that is stronger than the SecM(*Ec*). ⟨F⟩ (blue), $f_{fold}$ (orange), and $f_{\mathrm{FL}}$ (dashed orange) in dependence of $x_e$ ($K_p$ = 6.5 pN nm$^{-1}$, $\Delta G_{U \to F}$ = -20 kJ mol$^{-1}$, $x_c$ = 6.5 nm; $x^{‡}$ = 0.65 nm; Δt = 550 s) with $k_0$ = 3×10$^{-4}$ s$^{-1}$ (for SecM(*Ec*) AP) or $k_0$ = 6×10$^{-5}$ s$^{-1}$ (for a strong AP) as described by Eq. 1 – 8.

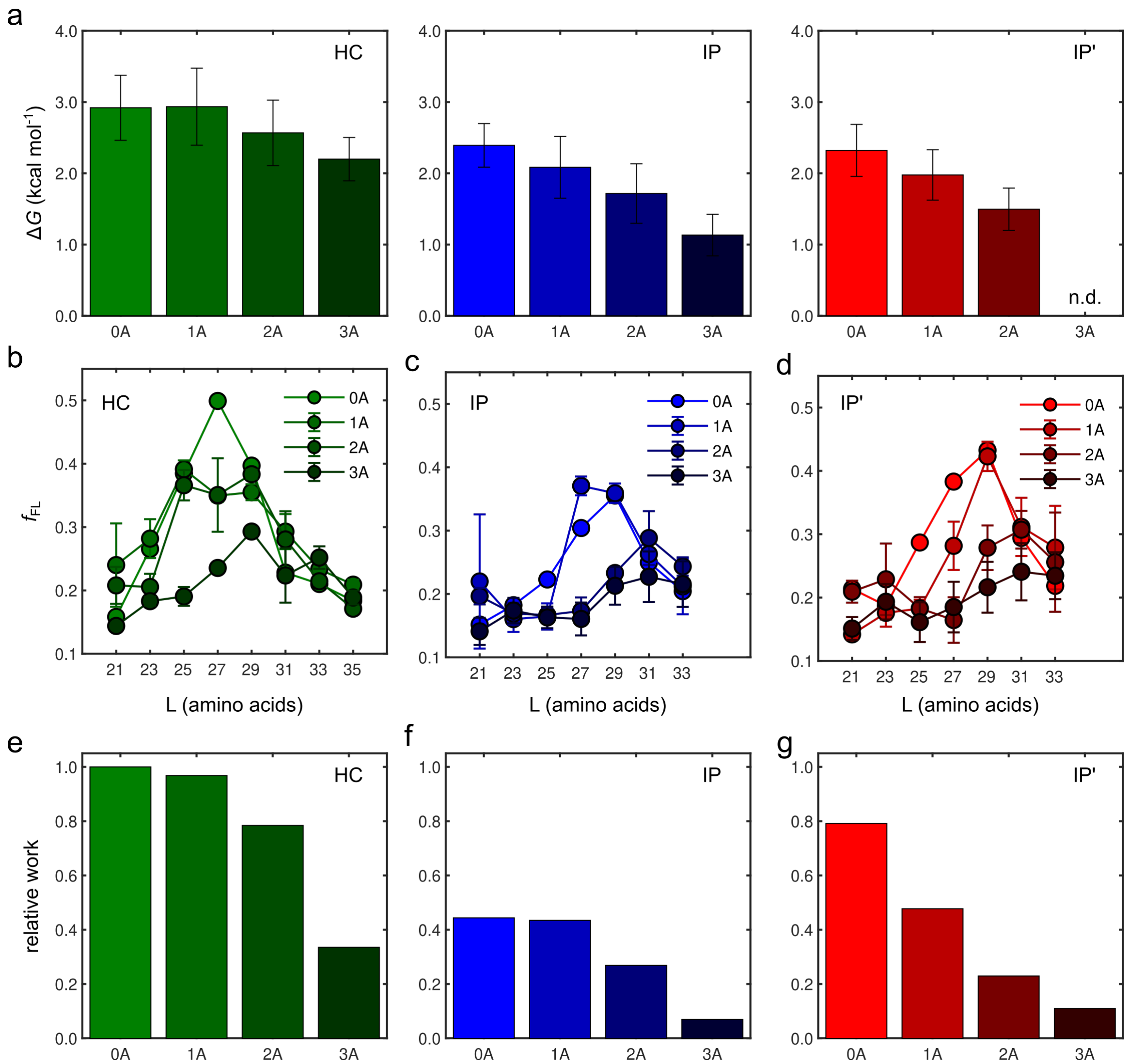


**Figure S4. The effect of decreasing hydrophobicity on the stability and folding of the protein *Maquettes*.** Gibbs free energy derived from CD chemical unfolding experiments for **(A)** HC, **(B)** IP and **(C)** IP’ with 0, 1 (L32A), 2 (L32A, I79A), or 3 (L32A, I79A, L87A) Ala substitutions. n.d.: not determined. **(B-G):** Force profile analysis for **(B)** HC, **(C)** IP and **(D)** IP’ with 0, 1, 2 or 3 Ala substitutions. Relative work exerted along the translated distance normalized to the HC data for (**E**) HC, (**F**) IP and (**G**) IP’ with 0, 1, 2 or 3 substitutions of bulky residues to alanine residues.

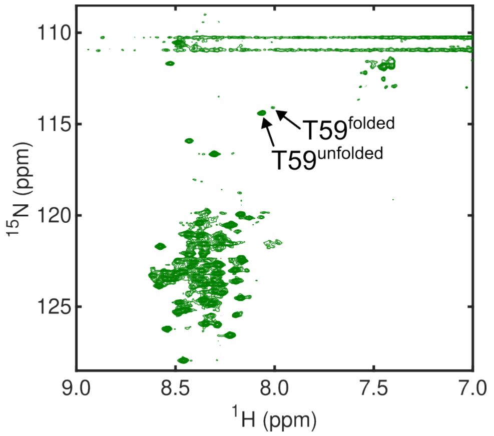


**Figure S5. $^{15}$N HSQC of HC at 10.5 M urea concentration.** Even at this concentration folded and unfolded conformation co-exist as exemplified for residue T59.

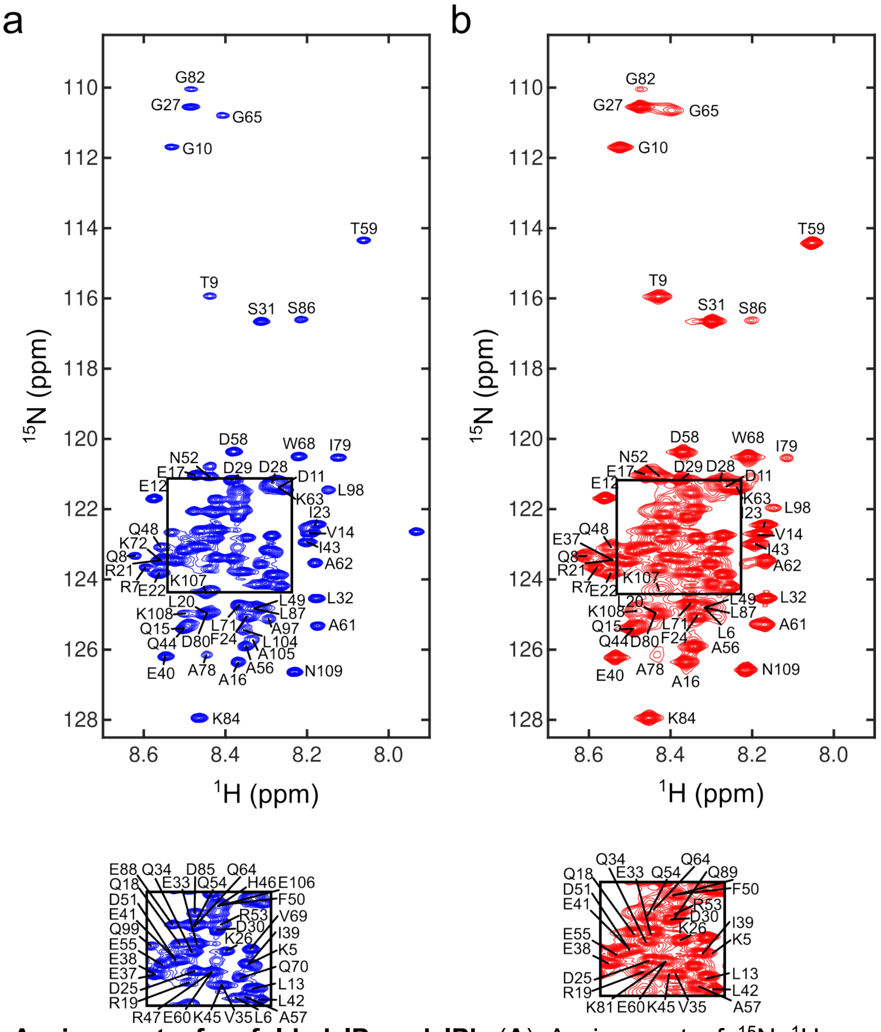


**Figure S6. Assignment of unfolded IP and IP'.** (**A**) Assignment of $^{15}$N–$^{1}$H resonances for unfolded IP and (**B**) assignment of $^{15}$N–$^{1}$H resonances for unfolded IP'.

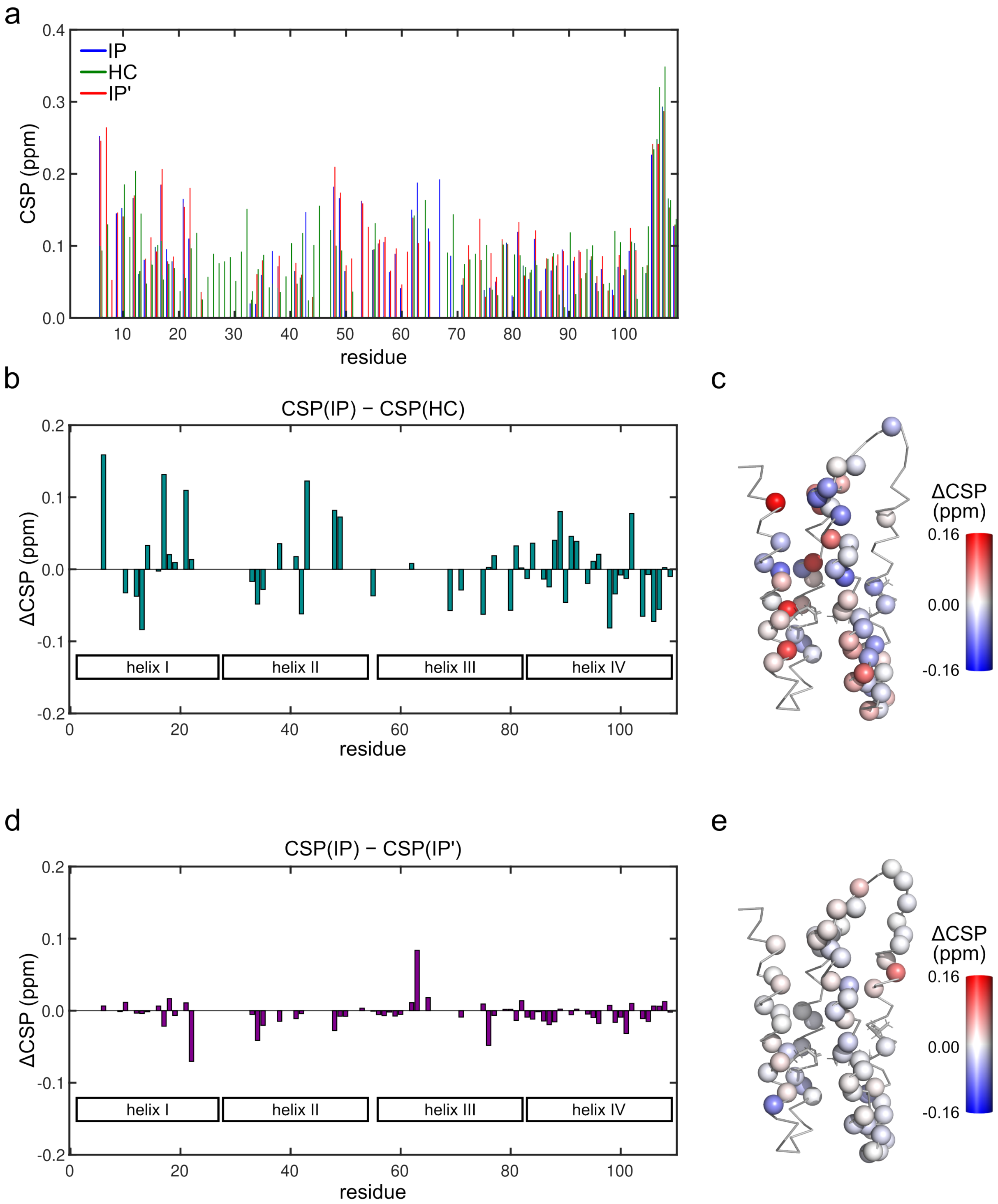


**Figure S7. Chemical shift perturbation of protein *Maquettes* during chemical unfolding experiments. (A)** CSPs of $^{15}$N-HSQC obtained for 0 M and 3 M of urea. **(B)** Differences in CSP between IP and HC (left, $\Delta CSP^{HC} = CSP^{IP} - CSP^{HC}$). **(C)** $\Delta CSP^{HC}$ mapped onto the NMR structure of IP (PDB: 6Z35). **(D)** Differences in CSP between IP and IP' ($\Delta CSP^{IP\text{-}IP'} = CSP^{IP} - CSP^{IP'}$). **(E)** $\Delta CSP^{IP'}$ mapped onto the NMR structure of IP (PDB: 6Z35). ΔCSP < 0 indicates residues in IP that are less surface-exposed than in HC/IP' and vice versa.

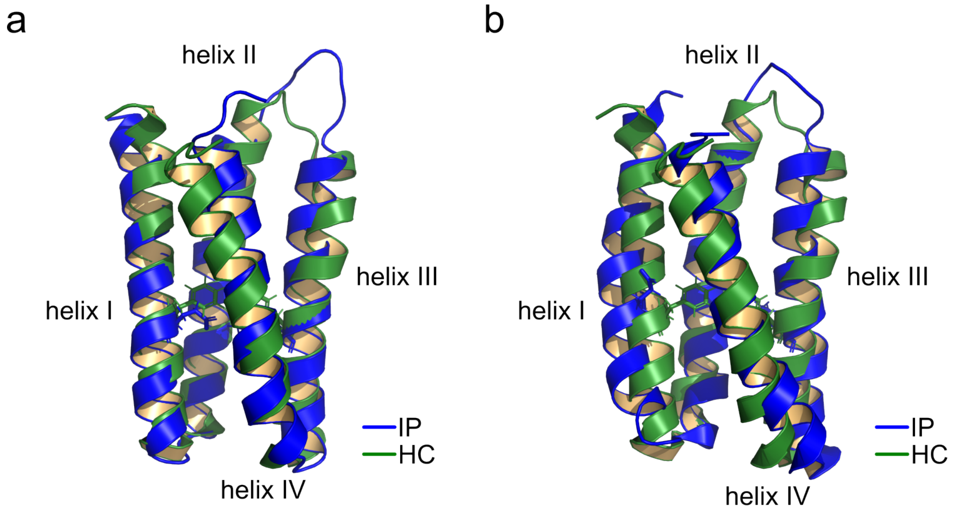


**Figure S8. Structural analysis of HC and IP**. Comparison of NMR structure of HC (green, PDB ID: 5TGW) with **(A)** IP (blue, PDB ID I: 6Z35) in the closed and **(B)** in the open conformation. F17/F72 (HC) and E17/K72 (IP) are shown as sticks.

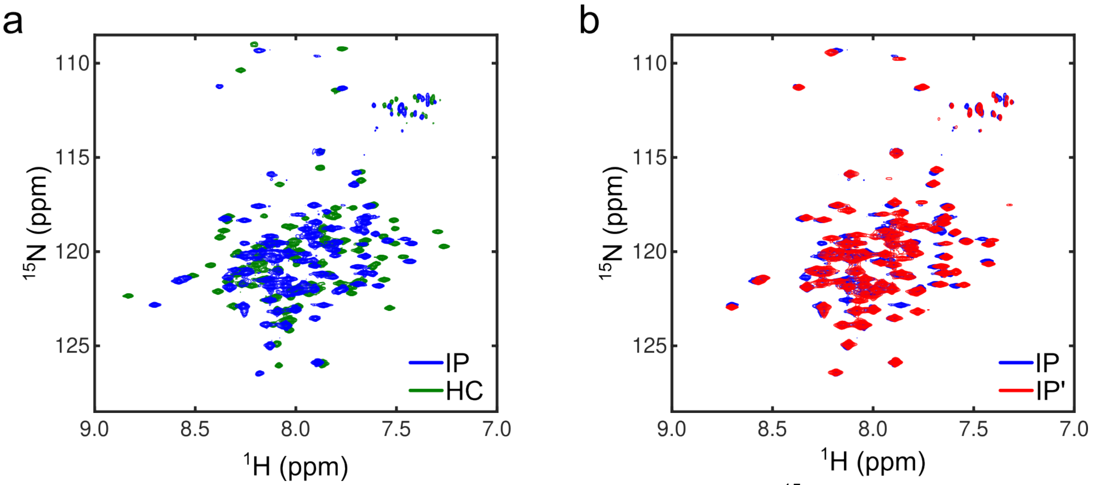


**Figure S9. HSCQ spectra of HC, IP, and IP'. (A)** Comparison of the $^{15}$N-HSQC of folded IP (blue) and folded HC (green). **(B)** Comparison of the $^{15}$N-HSQC of folded IP (blue) and folded IP' (red).

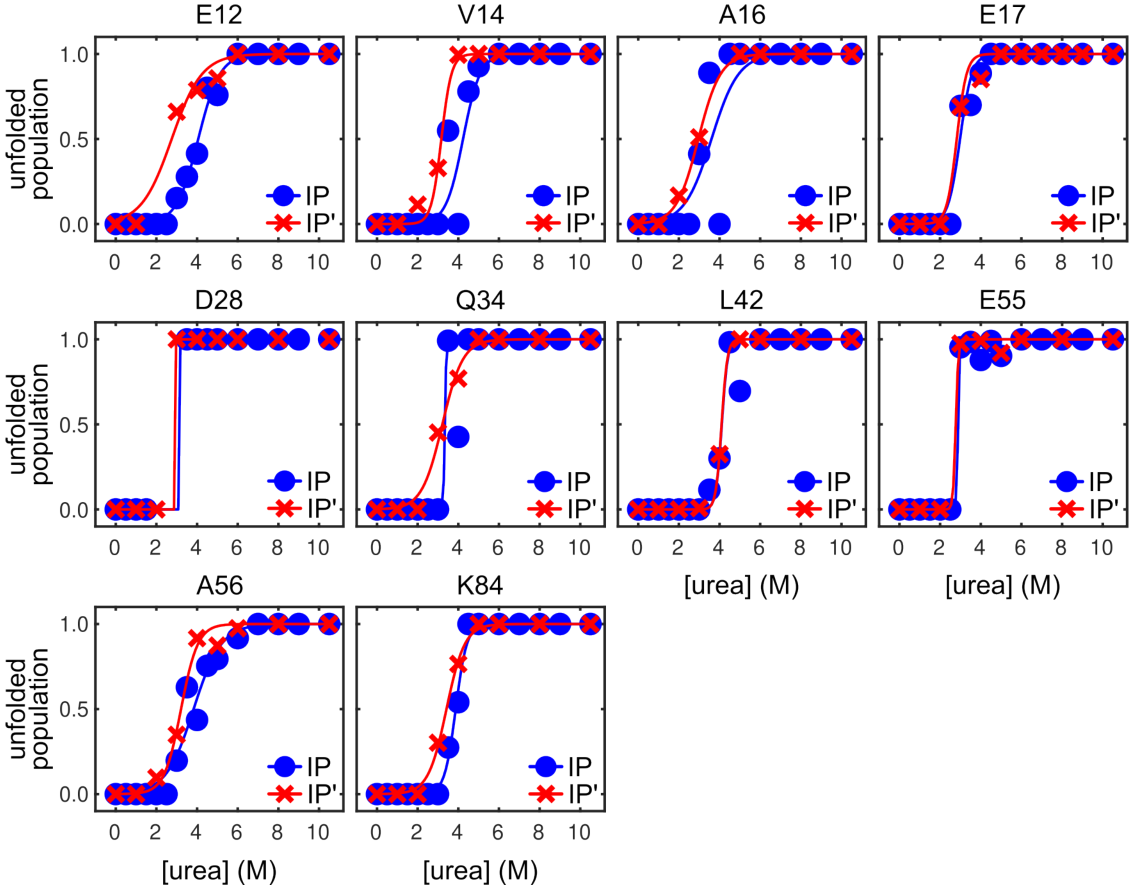


**Figure S10. Unfolding transitions for traceable residues in both IP and IP' investigated by NMR.** Eq. 9 is fitted to the data (see **Table S2**).

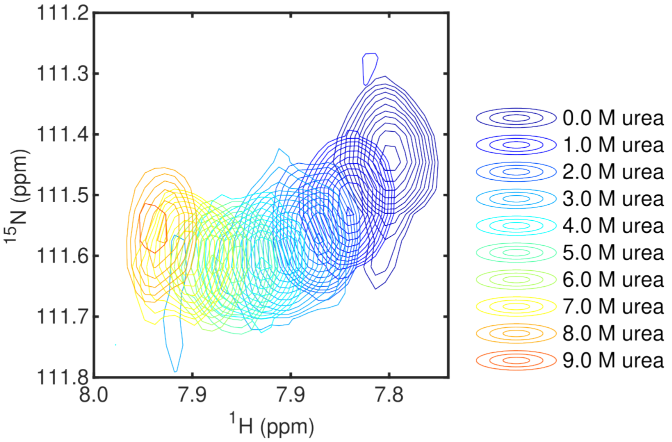


**Figure S11. G82 resonance of the folded conformation of HC at increasing urea concentrations.** The non-linear titration trajectory indicates an intermediate conformation that is in fast exchange with the folded conformation.

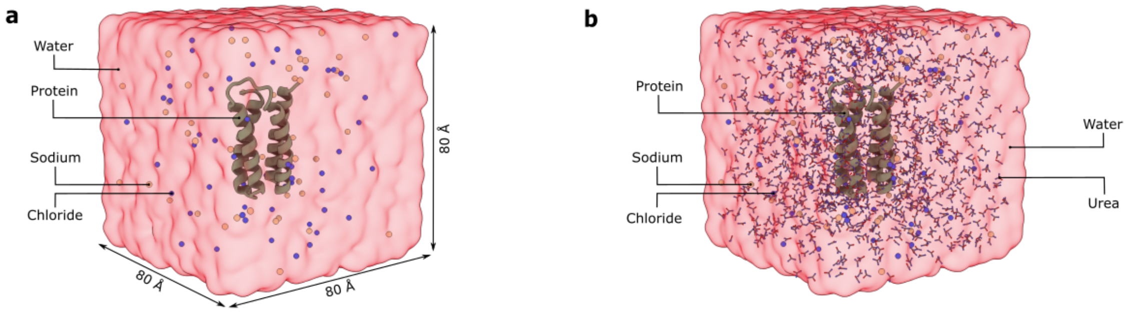


**Figure S12. Molecular dynamics simulation setup used for unfolding simulations. (A)** Representative simulation box containing the protein solvated in explicit water with $Na^+$ and $Cl^-$ ions in an ~80 × 80 × 80 $Å^3$ cubic box. **(B)** Representative simulation setup containing 2 M urea, showing the protein surrounded by explicit solvent, ions, and urea molecules. All systems were simulated under periodic boundary conditions. See Methods (main text) for further information.

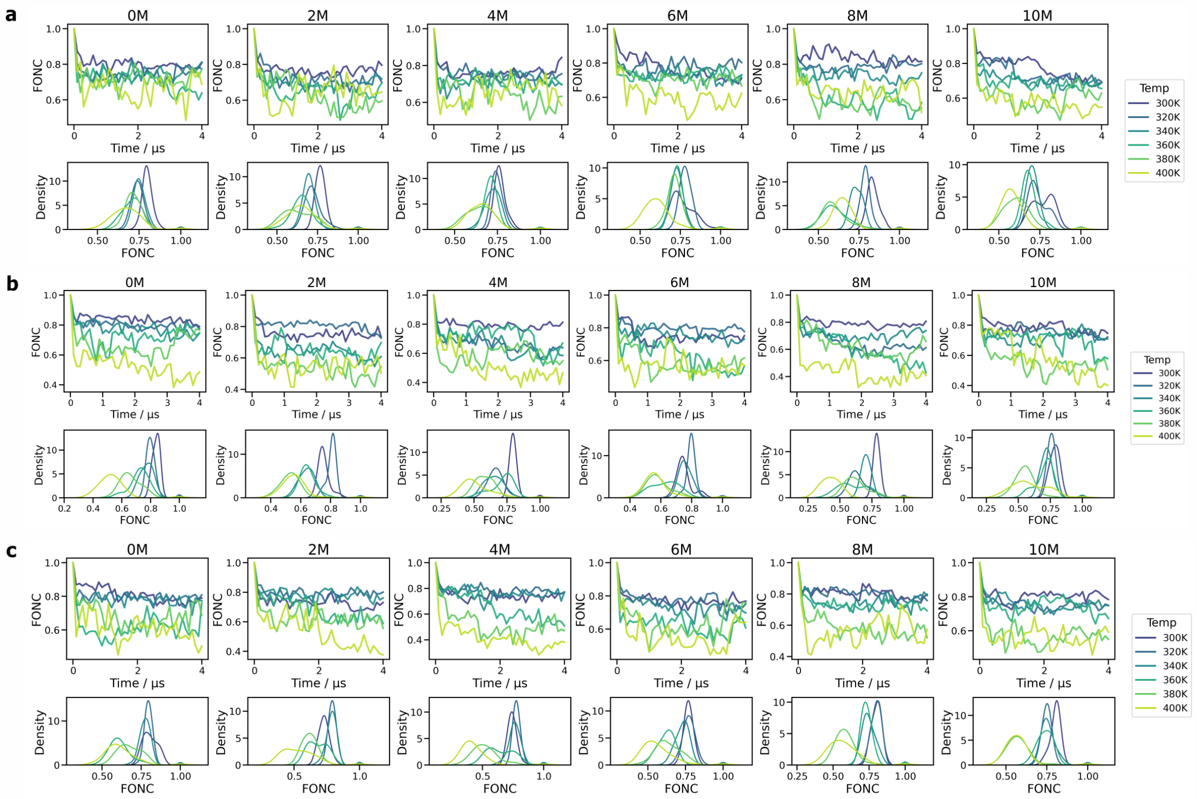


**Figure S13. Fraction of native contacts (FONC) across unfolding simulations under varying conditions.** Time evolution (top panels) and probability density distributions (bottom panels) of the fraction of native contacts are shown for **(A)** HC, **(B)** IP, and **(C)** IP' across urea concentrations ranging from 0–10 M and temperatures between 300–400 K. Each condition corresponds to a 4 μs simulation.

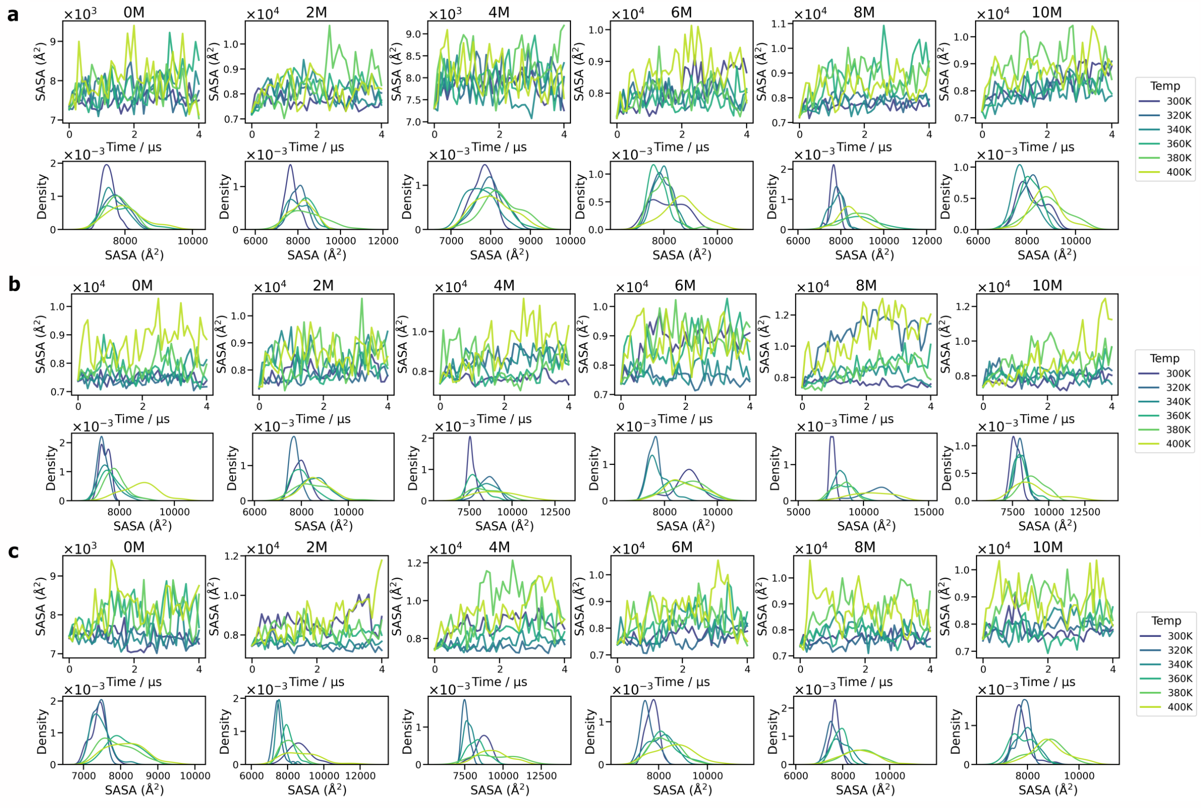


**Figure S14. Solvent-accessible surface area (SASA) across unfolding simulations under varying conditions.** Time evolution (top panels) and probability density distributions (bottom panels) of SASA are shown for **(A)** HC, **(B)** IP, and **(C)** IP' across urea concentrations ranging from 0–10 M and temperatures between 300–400 K.

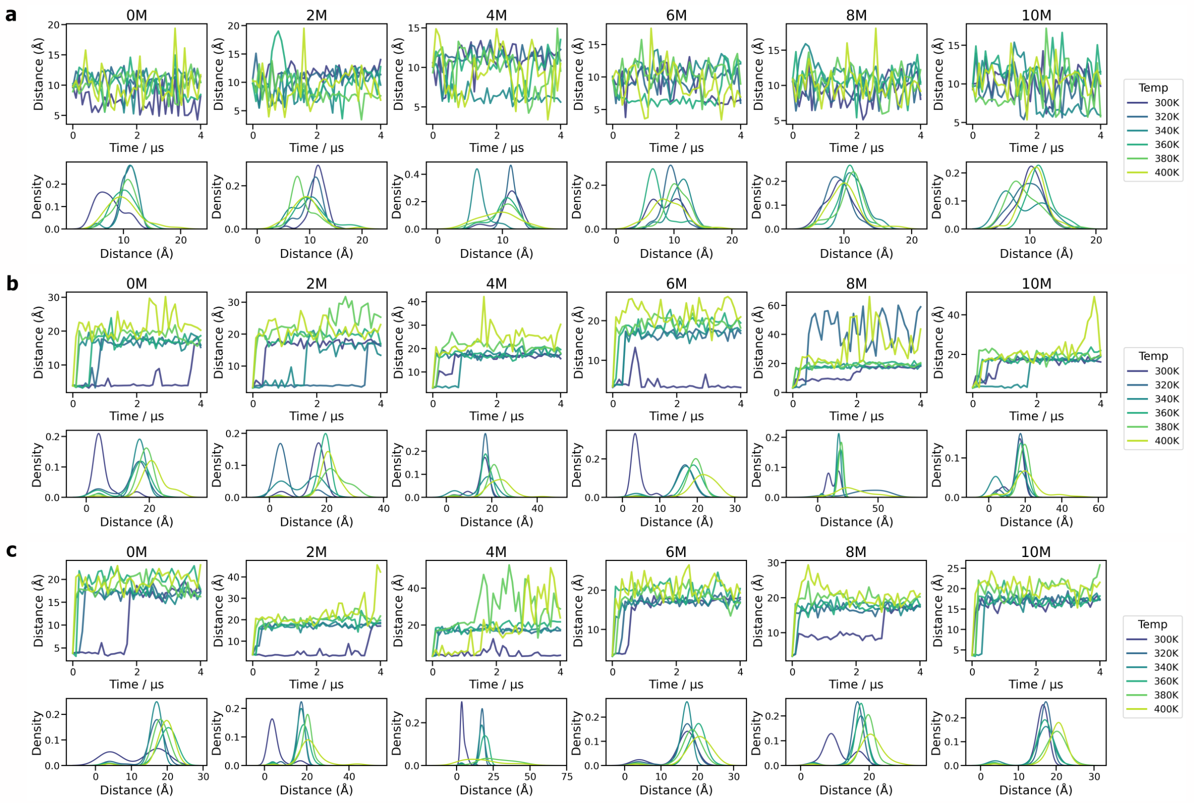


**Figure S15. Distance distributions monitoring core rearrangements during unfolding simulations.** Time evolution (top panels) and probability density distributions (bottom panels) of characteristic distances are shown for **(A)** HC, **(B)** IP, and **(C)** IP' across urea concentrations ranging from 0–10 M and temperatures between 300–400 K. Distances were measured between the terminal aromatic carbons of residues F17 and F72 in HC and between the carboxyl carbon of E17 and the terminal nitrogen atom of K72 in IP and IP'. Increasing denaturing strength shifts conformational populations toward larger distances.

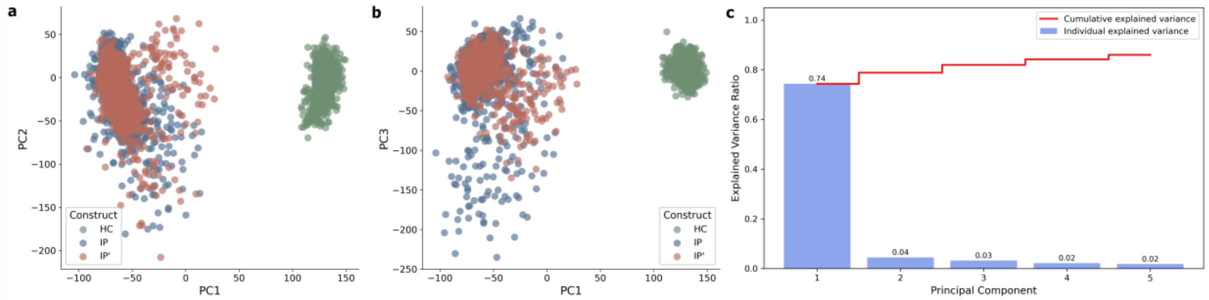


**Figure S16. Principal component analysis of conformational ensembles sampled during unfolding simulations.** Conformations from all simulations were projected into a PCA space using Cα coordinates. **(A)** Projection onto PC1–PC2, and **(B)** PC1–PC3 illustrates construct-dependent conformational sampling. HC occupies a distinct conformational space as compared to IP and IP', whereas the ion-pair-containing variants show substantial overlap. (**C**) Scree plot showing the variance explained by the first five principal components used for clustering analysis.

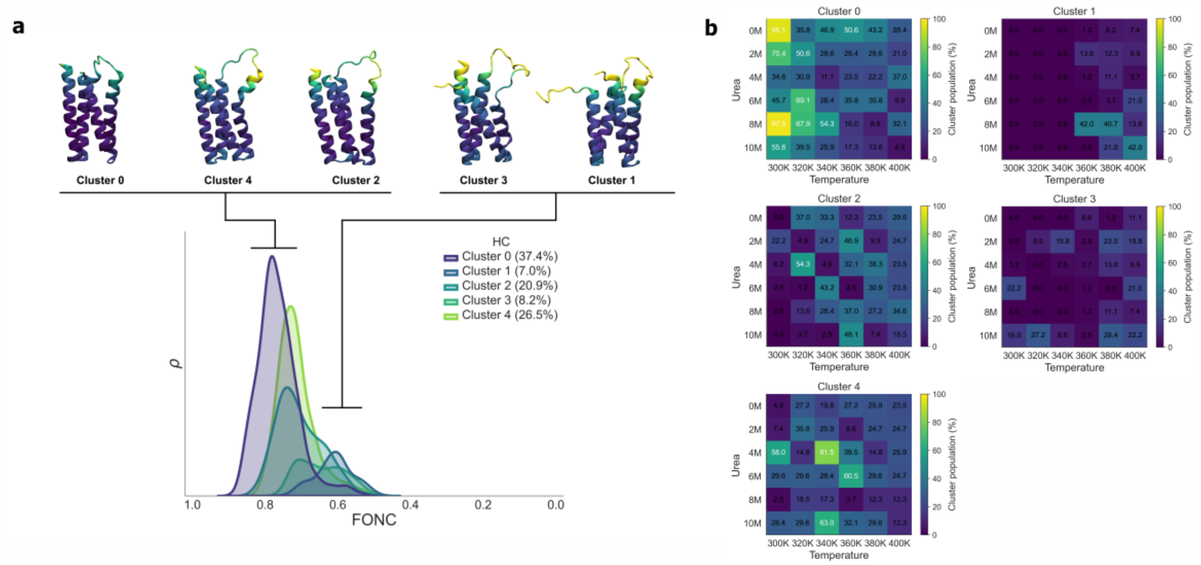


**Figure S17. PCA derived cluster analysis of HC unfolding intermediates. (A)** Representative structures for the five clusters identified from the combined simulation ensemble. Structures represent the conformer closest to the cluster-average structure by RMSD. Residues are colored by cluster-specific RMSF values to visualize structural variability. Corresponding FONC distributions and cluster populations are shown below. **(B)** Heatmaps showing the fraction of structures assigned to each cluster originating from different urea concentrations and temperatures, highlighting the simulation conditions that preferentially populate specific intermediates.

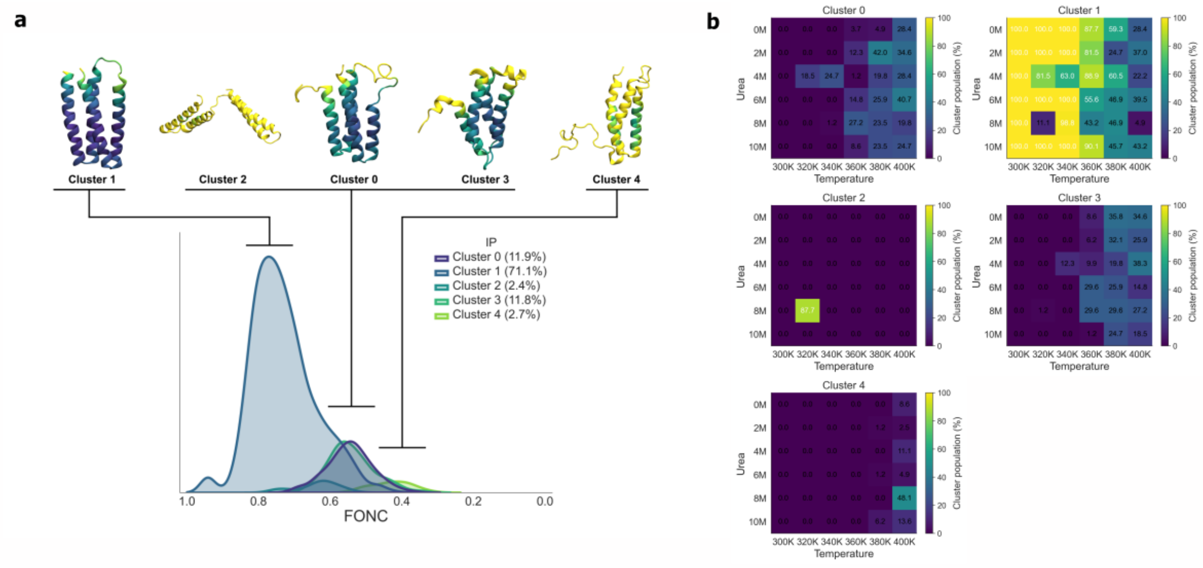


**Figure S18. PCA derived cluster analysis of IP unfolding intermediates. (A)** Representative structures for the five clusters identified from the combined simulation ensemble. Structures represent the conformer closest to the cluster-average structure by RMSD. Residues are colored by cluster-specific RMSF values to visualize structural variability. Corresponding FONC distributions and cluster populations are shown below. **(B)** Heatmaps showing the fraction of structures assigned to each cluster originating from different urea concentrations and temperatures, highlighting the simulation conditions that preferentially populate specific intermediates.

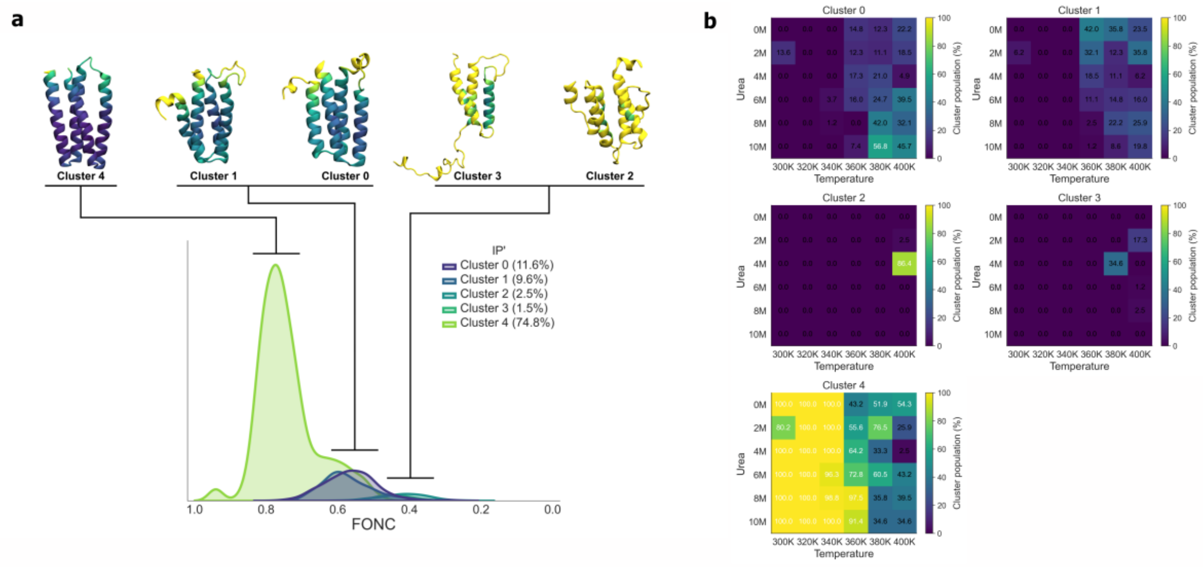


**Figure S19. PCA derived cluster analysis of IP' unfolding intermediates. (A)** Representative structures for the five clusters identified from the combined simulation ensemble. Structures represent the conformer closest to the cluster-average structure by RMSD. Residues are colored by cluster-specific RMSF values to visualize structural variability. Corresponding FONC distributions and cluster populations are shown below. **(B)** Heatmaps showing the fraction of structures assigned to each cluster originating from different urea concentrations and temperatures, highlighting the simulation conditions that preferentially populate specific intermediates.

**SI Tables**

**Table S1. Sequences of the Maquette constructs.** The ion-pair residues in IP and IP’ or equivalent Phe in HC are highlighted in red, the charge-shielding mutation N69 in IP’ and the equivalent V69 in IP and HC are highlighted in green and other residues mutated in this study are highlighted in bold.

| Construct | Sequence |
|---|---|
| IP | SEFEKLRQTG DELVQA**E**QRL REIFDKGDDD S**L**EQVLEEIE ELIQKHRQLF<br>DNRQEAADTE AAKQGDQW**V**Q L**K**QRFREA**I**D KGDKDS**L**EQL LEELEQALQK IRELAEKKN |
| IP’ | SEFEKLRQTG DELVQA**E**QRL REIFDKGDDD S**L**EQVLEEIE ELIQKHRQLF<br>DNRQEAADTE AAKQGDQW**N**Q L**K**QRFREA**I**D KGDKDS**L**EQL LEELEQALQK IRELAEKKN |
| HC | SEFEKLRQTG DELVQA**F**QRL REIFDKGDDD S**L**EQVLEEIE ELIQKHRQLF<br>DNRQEAADTE AAKQGDQW**V**Q L**F**QRFREA**I**D KGDKDS**L**EQL LEELEQALQK IRELAEKK |

**Table S2. Unfolding analysis of IP and IP'.** Fit values for unfolding analysis of IP and IP' (**Fig. S10**). Errors are given as ±1 SD.

| Residue | IP | | IP' | |
|---|---|---|---|---|
| | Growth rate ($M^{-1}$) | Midpoint transition (M) | Growth rate ($M^{-1}$) | Midpoint transition (M) |
| E12 | 0.0019 | 4.1 | 0.0013 | 2.8 |
| V14 | 0.0024 | 4.2 | 0.0040 | 3.2 |
| A16 | 0.0015 | 3.6 | 0.0018 | 3.0 |
| E17 | 0.0370 | 2.9 | 0.0170 | 2.7 |
| D28 | 0.3037 | 3.1 | 0.4927 | 2.9 |
| Q34 | 0.0300 | 3.3 | 0.0020 | 3.2 |
| L42 | 0.0070 | 4.1 | 0.0072 | 4.1 |
| E55 | 0.0264 | 2.9 | 0.0158 | 2.8 |
| A56 | 0.0015 | 3.8 | 0.0024 | 3.2 |
| K84 | 0.0037 | 3.9 | 0.0022 | 3.4 |
| Average | 0.04±0.09 | 3.6±0.5 | 0.05±0.15 | 3.1±0.4 |

**Table S3. Parameters for the global spectral dispersion analysis for IP, IP', and HC.**

| Sample | Growth rate ($10^{-3}$ ppm/$M^{-1}$) | Midpoint transition (M) | Amplitude (ppm) | Offset (ppm) |
|---|---|---|---|---|
| **IP** | -1.28 | 4.3 | 1.6 | 1.6 |
| **IP'** | -1.12 | 3.7 | 1.6 | 1.5 |
| **HC** | -1.65 | 8.2 | 1.9 | 1.5 |
